# Regulation of the formin INF2 by actin monomers and calcium-calmodulin

**DOI:** 10.1101/2025.10.06.680766

**Authors:** Miriam Lee, Aiman Jalmukhambetova, T. Emme Burgin, Henry N. Higgs

## Abstract

In response to increased intracellular calcium, the formin INF2 polymerizes 20-30% of the total cellular actin pool within 30 sec, suggesting robust regulation. INF2 regulation requires an auto-inhibitory interaction between the N-terminal Diaphanous Inhibitory Domain (DID) and the C-terminal Diaphanous Auto-regulatory Domain (DAD). DID mutations are dominantly linked to two human diseases, and constitutively activate INF2. However, DAD binding to actin monomers competes with DID binding, disrupting regulation. Here, we use a novel cell-free assay for detailed investigation of INF2 regulation. Contrary to our previous findings, INF2 inhibition does not require CAP proteins but does require actin ‘buffering’ by monomer-binding proteins such as profilin or thymosin. INF2 is activated by calcium-bound calmodulin (CALM) through CALM binding to the N-terminus. In addition, the N-terminus plays an important role in INF2 regulation beyond CALM binding. These findings support a role for actin monomer binding proteins in not only regulating overall actin dynamics but also in specific regulation of an actin polymerization factor.

**Summary:** In this work, a concerted regulatory mechanism for INF2 is described, in which INF2 is activated by a combination of calcium-calmodulin and free actin monomers. In other words, INF2 ‘senses’ actin monomers, making monomer binding proteins like profilin and thymosin important for INF2 regulation.

## Introduction

The actin cytoskeleton performs multiple roles in mammalian cells, including cell motility, endocytosis, membrane trafficking, cell adhesion, modulating mitochondrial function, and nuclear dynamics (Chhabra and Higgs, 2007; Blanchoin et al., 2014; Gurel et al., 2014b; Ulferts et al., 2021; Chakrabarti et al., 2021; Fung et al., 2023). The actin filaments used for these roles are largely generated from a common pool of actin monomers (Plastino and Blanchoin, 2018). Therefore, filament assembly and disassembly must be tightly regulated to ensure the appropriate actin-based structures assemble where and when needed. These actin-based structures are often transient, necessitating rapid filament turnover.

One dynamic actin-based system is a process we refer to as calcium-induced actin (CIA), which assembles in response to stimuli that raise cytoplasmic or nuclear calcium (Shao et al., 2015; Ji et al., 2015; Wales et al., 2016; Chakrabarti et al., 2018; Ulferts and Grosse, 2024). CIA can be exceptionally transient. In U2OS cells, peak actin polymerization occurs within 30 sec of stimulation by either ionomycin (a calcium ionophore) or histamine (a GPCR agonist), and returns to baseline within two minutes (Chakrabarti et al., 2018). Here, we show that CIA causes the assembly of a considerable percentage of the cell’s actin (40-60 million actin monomers) in this short period of time. The key actin polymerization factor in CIA is the formin INF2, which plays roles in multiple cellular processes, including mitochondrial dynamics (Korobova et al., 2013), vesicle trafficking (Madrid et al., 2010; Andrés-Delgado et al., 2010; Santos et al., 2020), organelle morphology (Ramabhadran et al., 2011; Tran et al., 2024), ER-organelle contacts (Chakrabarti et al., 2018; Schiavon et al., 2024), podosome dynamics (Panzer et al., 2016), and microtubule organization (Andrés-Delgado et al., 2012; Bartolini et al., 2016). Physiologically, INF2 is known to be important for glomerular function in the kidney, neuronal ischemic response, and placental development (Lamm et al., 2018; Calabrese et al., 2022; Labat-de-Hoz et al., 2024).

In view of CIA’s transience, tight regulation of INF2 is essential. A common regulatory mechanism for many formins is autoinhibition through the interaction between two domains: the Diaphanous Inhibitory Domain (DID, Li and Higgs, 2005) and the Diaphanous Auto-regulatory Domain (DAD, Alberts, 2001). In the formin mDia1, the DID and DAD interact with high affinity to inhibit the actin polymerization activity of the Formin Homology 2 (FH2) domain (Li and Higgs, 2003, 2005; Lammers et al., 2005). Activation of mDia1 occurs through GTP-bound RhoA binding to an N-terminal region overlapping the DID, disrupting DID/DAD interaction (Li and Higgs, 2003; Rose et al., 2005).

INF2 possesses both DID and DAD, and the DID/DAD interaction clearly plays a significant role in INF2 regulation. Point mutation of key residues in either the DID or DAD causes constitutive actin polymerization in cells (Chhabra et al., 2009; Ramabhadran et al., 2013). Furthermore, mutations in INF2’s DID are linked to two diseases, focal segmental glomerulosclerosis (FSGS, Brown et al., 2010) and Charcot-Marie Tooth disease (CMTD, Boyer et al., 2011). To date, mutations at >35 distinct positions in the DID have been identified in FSGS and CMTD patients (Labat-de-Hoz and Alonso, 2020). Where tested, these mutations cause constitutive INF2-mediated actin polymerization in cells (Bayraktar et al., 2020), consistent with their gain-of-function mechanism in disease progression (Subramanian et al., 2024).

Despite these findings, the regulatory mechanism for INF2 has been unclear. While INF2 is clearly tightly regulated in cells, purified full-length INF2 is constitutively active for actin assembly when mixed with actin monomers (Ramabhadran et al., 2013; A et al., 2019). The DID/DAD interaction for INF2 is >10-fold lower affinity than that of mDia1. Furthermore, actin monomers interact with the DAD in a manner that is competitive to DID binding (Ramabhadran et al., 2013). In past publications, we have put forward two possible mechanisms for INF2 regulation: 1) actin monomer binding proteins ‘buffer’ the actin/DAD interaction, allowing DID/DAD interaction and INF2 inhibition (Ramabhadran et al., 2013); and 2) a complex between acetylated actin and cyclase-associated proteins (CAPs) acts as a bridge between DID and DAD, resulting in INF2 inhibition (A et al., 2019, 2020).

The mechanism of INF2 activation has also been unclear, with several models being proposed (Labat-de-Hoz et al., 2024). One model, put forward by our laboratory, is that actin deacetylation triggers dissociation of the actin/CAP complex from INF2 (A et al., 2019, 2020). This model, however, does not provide a role for calcium in INF2 activation. A second model is that calcium-bound calmodulin (CALM) activates INF2 (Wales et al., 2016). Recently, a CALM binding site has been identified at the N-terminus of INF2, and mutations that eliminate CALM binding prevent cellular INF2 activation (Labat-de-Hoz et al., 2022).

In view of the relevance of INF2 regulation to human disease, resolution of the INF2 regulatory mechanism is necessary. In this paper, we develop a novel cell-free assay system to assess INF2 regulation. Our results support a role for actin monomer binding proteins (profilin and thymosin) in allowing INF2 auto-inhibition, with INF2 activation mediated by calcium-bound CALM interaction with the INF2 N-terminal region. Our results do not support a role for CAP proteins in INF2 regulation.

## Results

### Measuring the number of actin monomers polymerized during CIA

Calcium-induced actin (CIA) has been demonstrated in multiple publications (Shao et al., 2015; Ji et al., 2015; Wales et al., 2016; Chakrabarti et al., 2018). Two CIA characteristics are: 1) its rapidity, reaching a maximum within 1 min; and 2) its transience, returning to baseline within 2 min in cells (Chakrabarti et al., 2018). The amount of actin that is polymerized by CIA, however, has not been measured.

Here, we assessed the extent of CIA-mediated actin polymerization in U2-OS cells, using an assay in which actin filaments are separated from actin monomers by ultra-centrifugation, with actin filaments going to the pellet while actin monomers stay in the supernatant (Kage et al., 2022). In unstimulated cells, the percentage of polymerized actin varies in a range of 25-65% (mean of 44.8 ± 12.4%, **Fig. S1 A**). Stimulation with ionomycin causes a consistent increase in polymerized actin, with 31.6 ± 3.6% new actin polymerization occurring 30 sec after stimulation and returning to baseline within 2 min (**Fig. 1A, B. Fig. S1 B-E**). As shown previously (Chakrabarti et al., 2018), histamine causes more rapid and transient polymerization, reaching 20.7±1.8 % new actin polymerization in 10 sec and returning to baseline in 30 sec (**Fig. 1A, B. Fig. S1 F-H**). These dynamics largely mirror those measured using live-cell microscopy (**Fig. 1B,C**).

**Figure 1.**
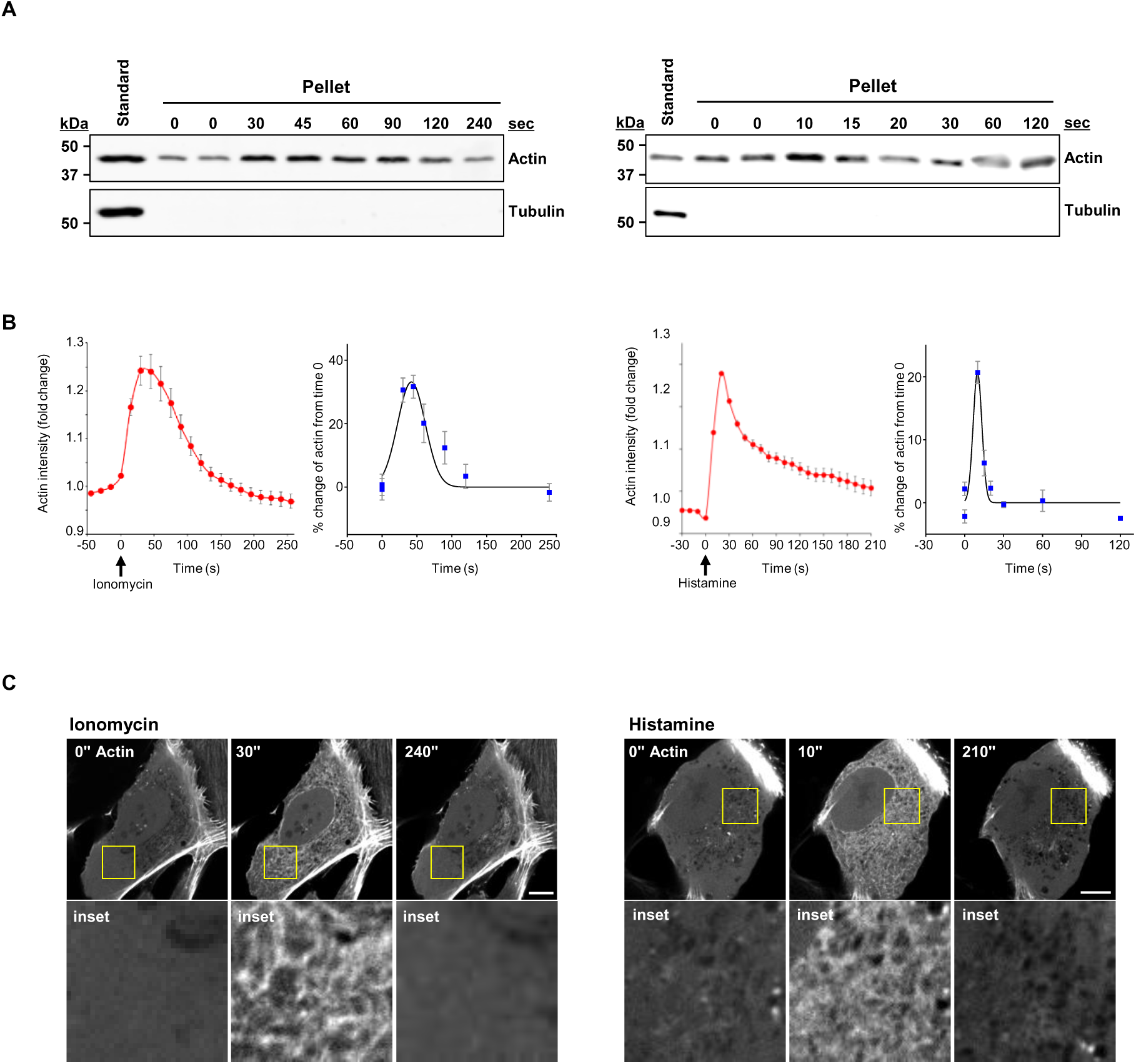
Calculating the amount of actin polymerized during CIA in U2-OS cells. **(A)** Representative western blot showing pelleted actin in U2-OS cells following ionomycin treatment (left) or histamine treatment (right) for the indicated times (sec). Tubulin also shown, with undetectable levels in the pellet. Standard, 40 ng of purified actin or tubulin. **(B)** Left: Time-course plots showing changes in cytoplasmic actin (GFP-F-tractin) in U2-OS cells and in pelleted actin following ionomycin addition. Fluorescence intensity and pelleted actin levels were each normalized to time 0 (F/F₀). GFP-F-tractin data represent n=33 cells, and actin pelleting data are from n=4 experiments; error bars indicate the standard error of the mean (SEM). Right: Same as left, but following histamine addition, with GFP-F-tractin data representing n=38 cells and actin pelleting data from n=3 experiments; error bars indicate SEM. **(C)** Time-lapse montage of an U2-OS cell transiently transfected with GFP-F-tractin, then stimulated with 4 μM ionomycin (left) or with 100 μM histamine (right) at time 0. Insets are magnified views of the boxed region. Scale bar: 10 μm; boxed region: 12 × 12 μm.

We used these results to calculate the number of actin molecules polymerized during CIA, using a cellular actin content of 192 million molecules/cell in U2-OS (the mean of three independent measurements of total actin content conducted in our laboratory (Hatch et al., 2016; A et al., 2020; Kage et al., 2022). Using an estimated cytoplasmic volume of 3.14 pL (A et al., 2020), the total U2-OS actin concentration is 101.6 μM. The 31.6% change in polymerized actin with ionomycin therefore represents 60.7 million actin monomers (32.1 μM), while the 20.7% change with histamine represents 39.7 million actin monomers (21.0 μM). The polymerization rate, therefore, is 2 million monomers/sec (1.1 μM/sec) for ionomycin and 3.97 million monomers/sec (2.1 μM/sec) for histamine. Considering that each actin monomer adds 2.7 nm to a filament, ionomycin and histamine polymerize 16.4 and 10.7 cm total actin filament length within 30 and 10 sec, respectively. Using our previously measured INF2 concentration in U2-OS cells (44,000 dimers/cell, A et al., 2020), we calculate that 1,380 and 902 actin monomers polymerize per INF2 dimer (∼4.3 μm filament per INF2 dimer) for ionomycin and histamine, respectively, if all INF2 dimers are active. The resulting rate of actin polymerization is 46 or 90 actin monomers per INF2 dimer per sec (respectively for ionomycin and histamine). While rapid, this rate is well below the maximum possible rate of INF2-mediated barbed end elongation, which we calculate to be 296 monomers/INF2 dimer/sec (see Materials and Methods for calculation). We point out that only a fraction of INF2 might be activated, so that the calculations of monomers per INF2 are theoretical.

In summary, this result demonstrates that CIA polymerizes a significant portion of the cell’s available actin monomers in seconds, with depolymerization of this actin occurring almost as quickly. This level of control requires tight but readily reversible regulation of INF2 activity. We sought to understand the rapid regulation of INF2 in more detail.

### Development of a cell-free assay for INF2 activity

In cells, the time course of CIA follows closely behind the time course of calcium increase and decrease (Chakrabarti et al., 2018), suggesting tightly coupled regulation. Despite INF2’s tight regulation in cells, purified full-length INF2 constitutively stimulates actin polymerization, with similar efficiency to a construct lacking the DID (called INF2-FFC, **Fig. 2 A**) in pyrene-actin polymerization assays containing only INF2 and actin (**Fig. S2 A**). This discrepancy in INF2 regulation prompted us to develop a cell-free assay that replicates the calcium-dependent regulation of INF2 in cells, but also allows addition of soluble components. Our system utilizes INF2-knockout (KO) U2-OS cells which stably express GFP-INF2-CAAX (**Fig. 2 B**). Consistent with previous studies, GFP-INF2-CAAX enriches on the ER, and histamine stimulation leads to rapid and transient actin polymerization in the GFP-INF2-CAAX cell line, but not in the control line expressing GFP alone (**Fig. 2 C).**

**Figure 2.**
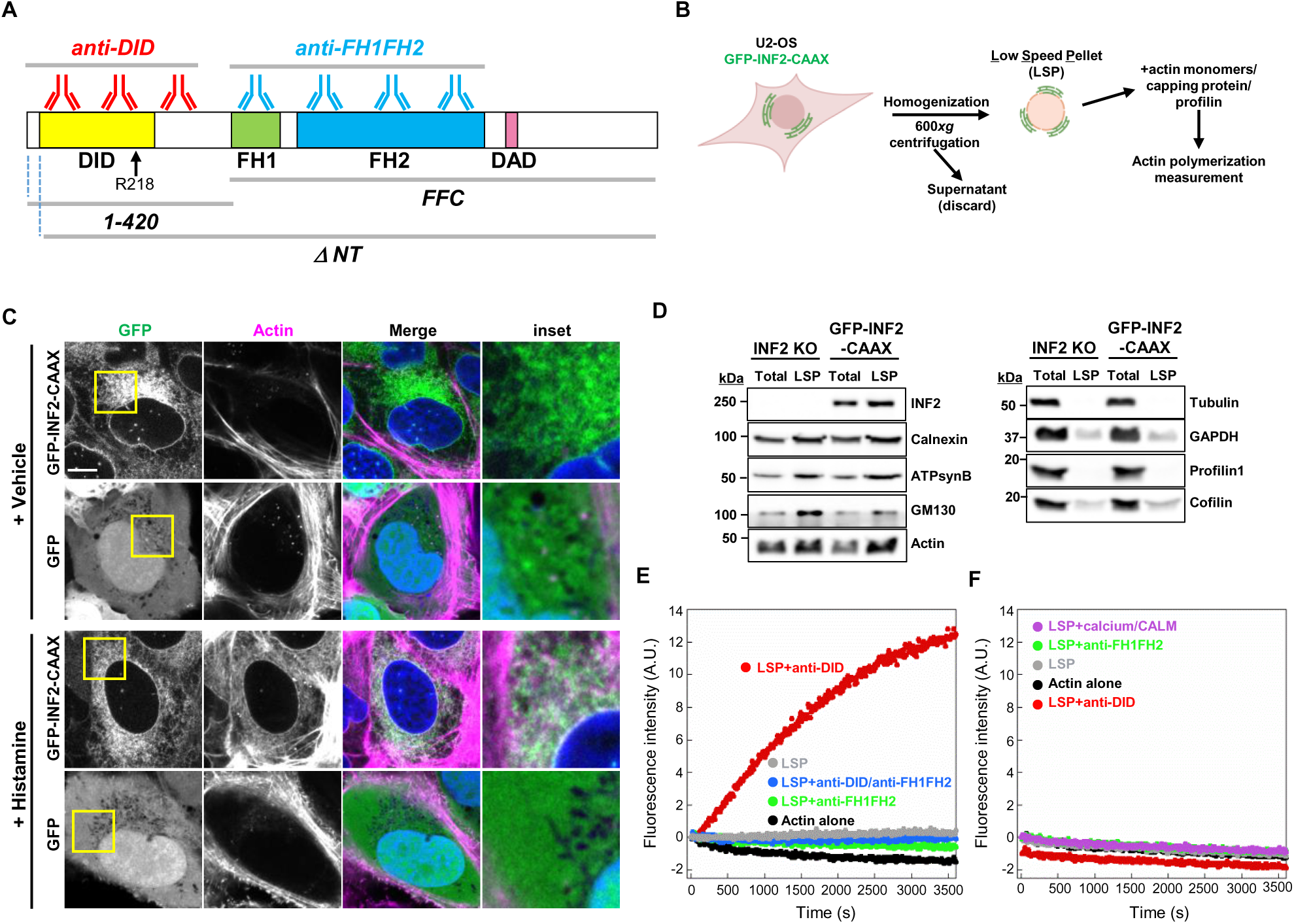
A cell-free assay for INF2-mediated actin polymerization. **(A)** Schematic diagram of the INF2 domains, highlighting the regions targeted by polyclonal antibodies, the disease-associated residue R218, as well as the 1-420, ΔNT, and FFC constructs used in this study. Domain boundaries (human INF2): N-terminus, 1-34; DID, 35-259; FH1, 421-520; FH2, 535-941; DAD, 967-995. **(B)** Schematic of the cell-free assay. U2-OS INF2 KO cells stably expressing GFP-INF2-CAAX are homogenized, followed by low speed centrifugation (600xg), after which the pellet (LSP) is used. Actin polymerization activity of the LSP is measured using pyrene actin assay, in the presence of actin monomer (2 μM, 5% pyrene-labeled), capping protein (50 nM) and profilin (6 μM). **(C)** U2-OS INF2 KO cells stably expressing GFP-INF2-CAAX or GFP were treated with vehicle or 100 μM histamine for 30 seconds, then fixed and stained with TRITC-phalloidin (to visualize actin filaments) and DAPI (to stain the nucleus). Scale bar: 10 μm; boxed region: 12 × 12 μm. **(D)** Western blots of total cell lysate and LSP from either U2-OS INF2 KO cells or GFP-INF2-CAAX stable cells. Equal protein (5 μg) was loaded in each lane and blot was probed with the indicated antibodies. **(E, F)** Representative pyrene-actin polymerization assays using LSP from GFP-INF2-CAAX stable cells (E) or from INF2 KO cells (F). Each reaction contains 10 μg LSP protein, 2 μM added actin monomer (5% pyrene-labeled), 50 nM capping protein, and 6 μM profilin, with or without the indicated components (167 nM anti-DID antibody, 167 nM anti-FH1FH2 antibody, 1 μM calcium with 250 nM CALM, as specified). Experiments were performed in triplicate with similar results, and the data shown are a representative experiment.

Using this cell line, we developed a simple and rapid method for enriching ER membrane (**Fig. 2 B**). Cells are homogenized by dounce in detergent-free actin polymerization buffer, then subjected to low-speed centrifugation (600xg). The resulting pellet (referred to as the LSP, for low-speed pellet) contains nuclei and is also enriched for markers of ER, mitochondria and Golgi, as well as for GFP-INF2-CAAX, but is depleted for cytoplasmic proteins such as tubulin, GAPDH, profilin and cofilin, which remain in the supernatant after cell lysis (**Fig 2 D**). We tested enrichment of other actin-binding proteins and found that mDia1, myosin II heavy chain, Arp2/3 complex, CAP proteins, and fascin are depleted in LSP, whereas mDia2, FMNL formins, and alpha-actinins are enriched (**Fig. S2 B**).

Actin is also enriched in the LSP (**Fig. 2 D**), and we wished to characterize this actin before using the system to study INF2. LSP actin accounts for 51±5.8% of the total cellular actin **(Fig. S2 C)**. By fluorescence microscopy of LSP, there is TRITC-phalloidin labeling near to the DAPI label from both GFP-INF2-CAAX and the GFP alone-expressing cells (**Fig. S2 D**), suggesting that much of the LSP actin may be polymerized in an INF2-independent manner. Using the afore-mentioned pelleting assay to assess the polymerization state of LSP actin, >95% of the actin is in the detergent-extracted pellet fraction for both INF2-CAAX LSP and GFP LSP. Overnight treatment of LSP with the actin-sequestering molecule latrunculin A (LatA) prior to addition of phalloidin changes this value to 55.5% **(Fig. S2 E and F)**, suggesting that about half of the LSP actin is stably polymerized while half is capable of depolymerization. In subsequent actin polymerization assays, the total LSP actin is present at 2.2 μM (see Methods).

We tested the ability of LSP to stimulate polymerization of added actin monomers, using pyrene-actin fluorescence (2 μM actin monomers, 5% pyrene-labeled). To suppress background actin polymerization, we also added 6 μM profilin and 50 nM capping protein. Profilin inhibits spontaneous actin nucleation and pointed end elongation, while capping protein inhibits barbed end polymerization (Pollard et al., 2000). The combination of profilin and capping protein prevents actin polymerization over the 1 hr assay period (**Fig. 2 E**) or overnight (**Fig. S2 G**). Inclusion of LSP from GFP-INF2-CAAX cells does not stimulate actin polymerization (**Fig. 2 E, Fig. S2 G**), suggesting that the actin already present in the LSP does not contribute to polymerization in the absence of stimulatory factors.

We then attempted to manipulate INF2-mediated actin polymerization using two different polyclonal antibodies, hypothesizing that these antibodies might disrupt the functions of their targets. One antibody binds the INF2 DID-containing N-terminal region (anti-DID), and the other binds the INF2 FH1-FH2 region (anti-FH1FH2) (**Fig. 2 A**). Treatment of LSP with anti-DID alone causes significant actin polymerization, while addition of anti-FH1-FH2 inhibits the polymerization caused by anti-DID (**Fig. 2 E**). LSP prepared from the INF2 KO cell line displays no actin polymerization activity in the presence of anti-DID (**Fig. 2 F**). These results suggest that our cell-free system is capable of assessing INF2 activity, with anti-DID activating actin polymerization from the LSP fraction by disrupting INF2 inhibition, while anti-FH1FH2 blocks polymerization by inhibiting INF2’s interaction with actin.

### Calcium/calmodulin activates INF2 in cell-free assays

In cells, INF2 is activated by calcium (Shao et al., 2015; Wales et al., 2016; Chakrabarti et al., 2018; Labat-de-Hoz et al., 2022), with calmodulin (CALM) appearing to mediate this activation (Wales et al., 2016; Labat-de-Hoz et al., 2022). We tested the effects of calcium and CALM in our LSP assay. Addition of calcium alone or CALM alone has a slight stimulatory effect on actin polymerization, while the combination of calcium and CALM triggers robust actin polymerization (**Fig. 3 A**). The effect of calcium alone may be due to calmodulin present in the LSP preparation **(Fig. S2 B)**. Supporting this conclusion, actin polymerization caused by either calcium alone or by calcium with calmodulin is blocked by the calmodulin antagonist W-7 **(Fig. S2 H)**. Titration of CALM in the presence of 1 μM free calcium reveals an EC50 of 9.1 nM for CALM (**Fig. 3 B**), whereas titration of calcium in the presence of 1 μM CALM shows a sigmoidal response with an EC50 of 240 nM and a Hill coefficient of 8.9 for free calcium (**Fig. 3 C**), likely reflecting cooperative binding to the four-calcium binding sites present in calmodulin. Calcium/CALM-mediated activation is inhibited by anti-FH1FH2 (**Fig. 3 D**). LSP from INF2 KO cells does not respond to calcium/CALM treatment (**Fig. 2 F**).

**Figure 3.**
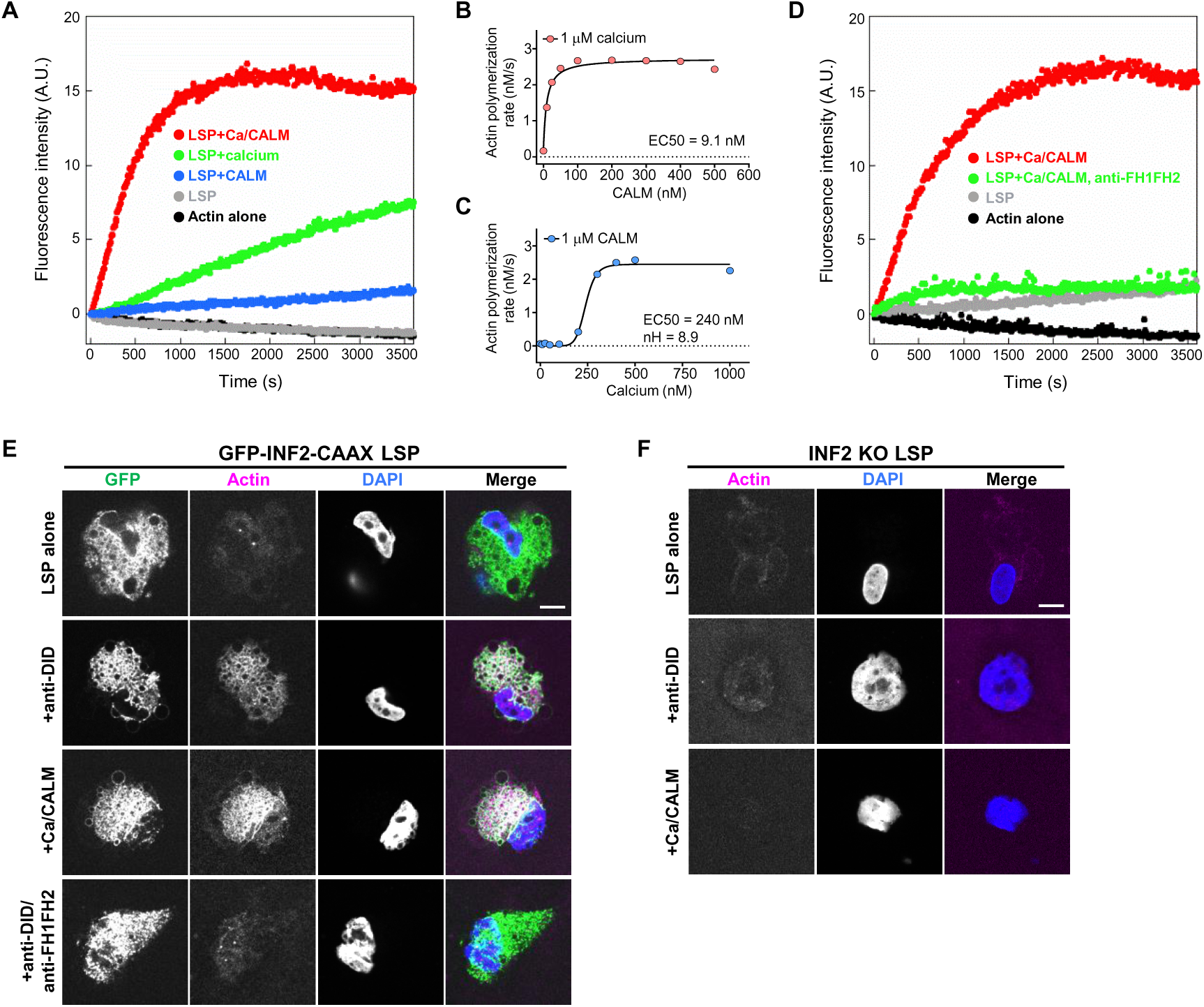
Calcium/calmodulin activates INF2-mediated actin polymerization in a cell-free assay system. **(A)** Pyrene-actin polymerization assay containing 2 μM actin (5% pyrene-labeled), 50 nM capping protein, 6 μM profilin, and 10 μg LSP from GFP-INF2-CAAX stable cells, with or without 1 μM calcium and 1 μM CALM. Experiments were performed in triplicate with similar results, and the data shown are a representative experiment. **(B, C)** Titration curves for CALM in the presence of 1 μM calcium (B) and free calcium in the presence of 1 μM CALM (C). Polymerization rates were calculated from the linear phase of the pyrene-actin assay (0-500 seconds). *nH*= Hill coefficient. **(D)** Pyrene-actin polymerization assay performed as in (A), with the indicated components (1 μM calcium, 250 nM CALM and/or 670 nM anti-FH1FH2 antibody). Experiments were performed in triplicate with similar results, and the data shown are a representative experiment. **(E, F)** Representative confocal microscopy images showing polymerized actin around LSP from GFP-INF2-CAAX stable cells (E) or INF2 KO cells (F). Each reaction contains 10 μg LSP, 2 μM added actin monomer (10% TAMRA-labeled), 50 nM capping protein, and 6 μM profilin, with or without the indicated components (167 nM anti-DID antibody, 167 nM anti-FH1FH2 antibody, 1 μM calcium with 100 nM CALM, as specified).

We also used fluorescence microscopy with TAMRA-labeled actin monomers, to examine incorporation of new actin monomers into the ER-enriched membranes (as opposed to TRITC-phalloidin labeling (**Fig. S2D**), which stains both new and existing filaments in LSP). Similar to the pyrene-actin assays, anti-DID or calcium/CALM stimulates actin polymerization around nuclei in GFP-INF2-CAAX expressing cells (**Fig. 3 E**). In contrast, control INF2 KO cells display no TAMRA-actin accumulation around nuclei (**Fig. 3 F**). These results show that addition of physiologically-relevant calcium concentrations along with nanomolar concentrations of CALM are sufficient to activate INF2.

### The FSGS-linked R218Q mutant of INF2 causes constitutive polymerization in cell-free assays

Disease-linked INF2 mutations in the DID region have been shown to cause constitutive INF2 activity (Bayraktar et al., 2020; A et al., 2019), and one of the most common of these mutations is R218Q (**Fig. 2 A**). In intact U2-OS cells, transient expression of GFP-INF2-CAAX-R218Q causes constitutive actin polymerization, as opposed to GFP-INF2-CAAX-WT, which displays histamine-sensitive actin polymerization (**Fig. 4, A and B**). We tested these constructs in our cell-free assay. Compared to our stable expression system (**Fig. 2**), LSP from cells transiently expressing GFP-INF2-CAAX-WT display a background level of actin polymerization in the absence of stimulus (**Fig. 4 C**). However, actin polymerization is substantially activated by anti-DID or calcium/CALM treatment (**Fig. 4 C**). In contrast, LSP from GFP-INF2-CAAX-R218Q cells displays maximal actin polymerization that is not further stimulated by calcium/CALM and is even somewhat decreased by anti-DID (**Fig. 4 D**). These results suggest that the R218Q mutation disrupts INF2 regulation, rendering it constitutively active.

**Figure 4.**
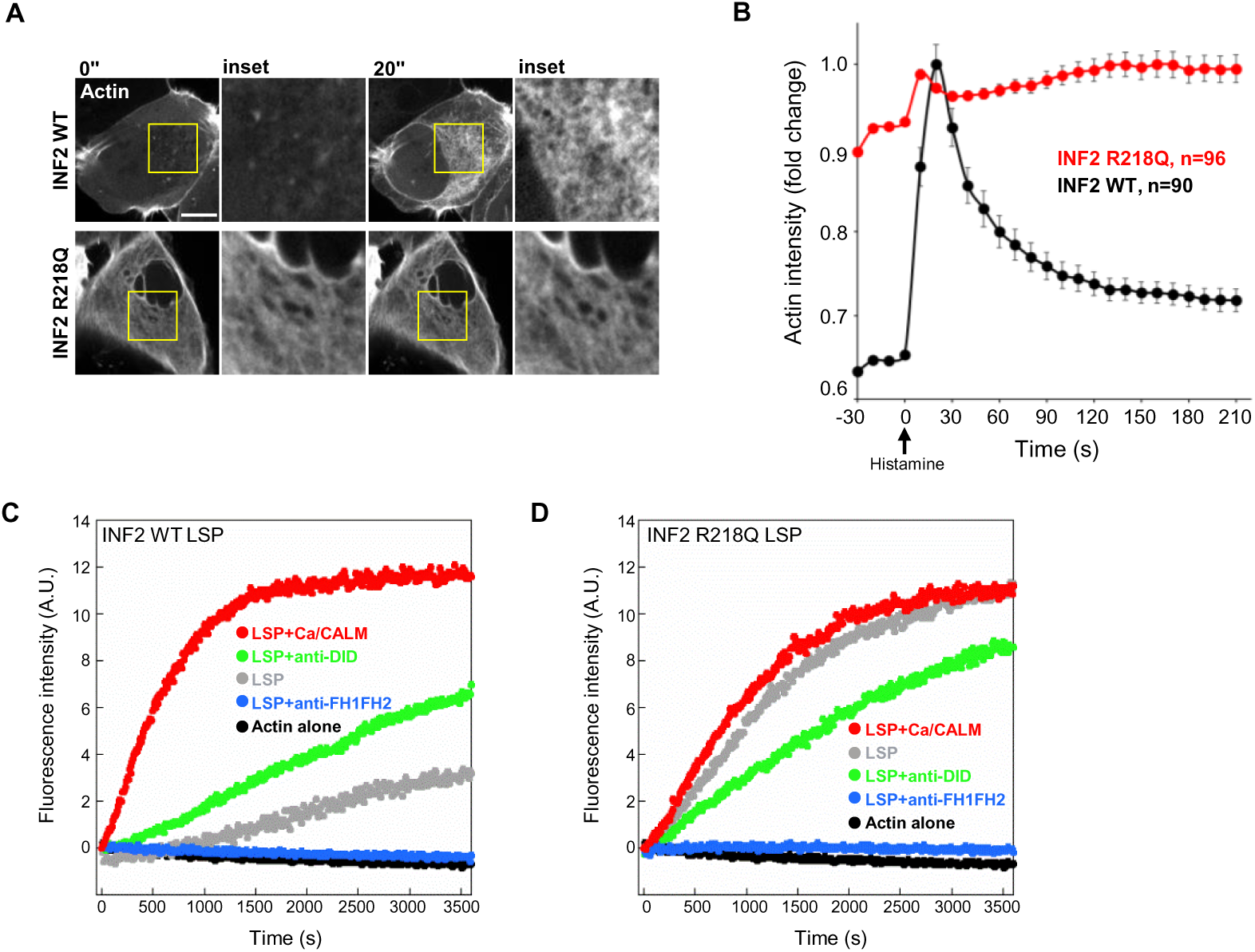
The FSGS-linked INF2 R218Q mutant exhibits constitutive activity in both intact cells and cell-free assays. **(A)** Time-lapse montage of U2-OS INF2 KO cells transiently expressing mApple-F-tractin along with either GFP-INF2 wild-type or GFP-INF2 R218Q, stimulated with 100 μM histamine at time 0. Insets show magnified views of the boxed regions. Scale bar: 10 μm; boxed region: 12 × 12 μm. **(B)** Time-course plot showing changes in cytoplasmic actin (mApple-F-tractin) following histamine addition. Y-axis values represent fluorescence intensity at each time point, normalized to the maximum fluorescence intensity observed (20 sec time point in INF2 WT). Error bars represent the standard error of the mean (SEM) of the indicated cell numbers. **(C, D)** Pyrene-actin polymerization assays containing 2 μM actin (5% pyrene-labeled), 50 nM capping protein, 6 μM profilin, and 10 μg of LSP from U2-OS INF2 KO cells transiently expressing either GFP-INF2-CAAX WT (C) or GFP-INF2-CAAX R218Q (D), with the indicated components (330 nM anti-DID antibody, 209 nM anti-FH1FH2 antibody, or 1 μM calcium with 100 nM CALM). Experiments were performed in triplicate with similar results, and the data shown are a representative experiment.

Collectively, these findings suggest that our cell-free assay recapitulates regulated INF2-mediated actin polymerization. Actin polymerization is activated by either anti-INF2-DID or by calcium/CALM, with anti-INF2-FH1FH2 antibody inhibiting actin polymerization. Neither anti-DID nor calcium/CALM treatment activates actin polymerization in INF2 KO extract. Similar to intact cells, the FSGS-linked R218Q mutant causes constitutive actin polymerization. In the next section, we use this system to test the effect of CAP proteins on INF2 activity.

### CAP proteins are not essential for INF2 regulation

Previously, we found evidence that CAP proteins participate in INF2 regulation (A et al., 2019, 2020). Surprisingly, however, both CAP1 and CAP2 are at low levels in LSP (**Fig. S2 B**), despite tight INF2 regulation. To test a role for CAP proteins directly, we depleted both CAP1 and CAP2 by siRNA and measured INF2 activity in the cell-free assay. siRNA treatment reduces CAP1 and CAP2 levels significantly in both the total cell lysate and the LSP fraction, while having no apparent effect on the levels of INF2 or actin (**Fig. 5 A**). The LSP from CAP1/2 KD cells behaves similarly to LSP from control cells, requiring activation by either anti-DID or calcium/CALM to induce actin polymerization (**Fig. 5, B and C**). In addition, supplementing the LSP with purified CAP2 has minimal effect on INF2 activity (**Fig. 5 D**), suggesting that CAPs are not required to keep INF2 inactive in the cell-free assay.

**Figure 5.**
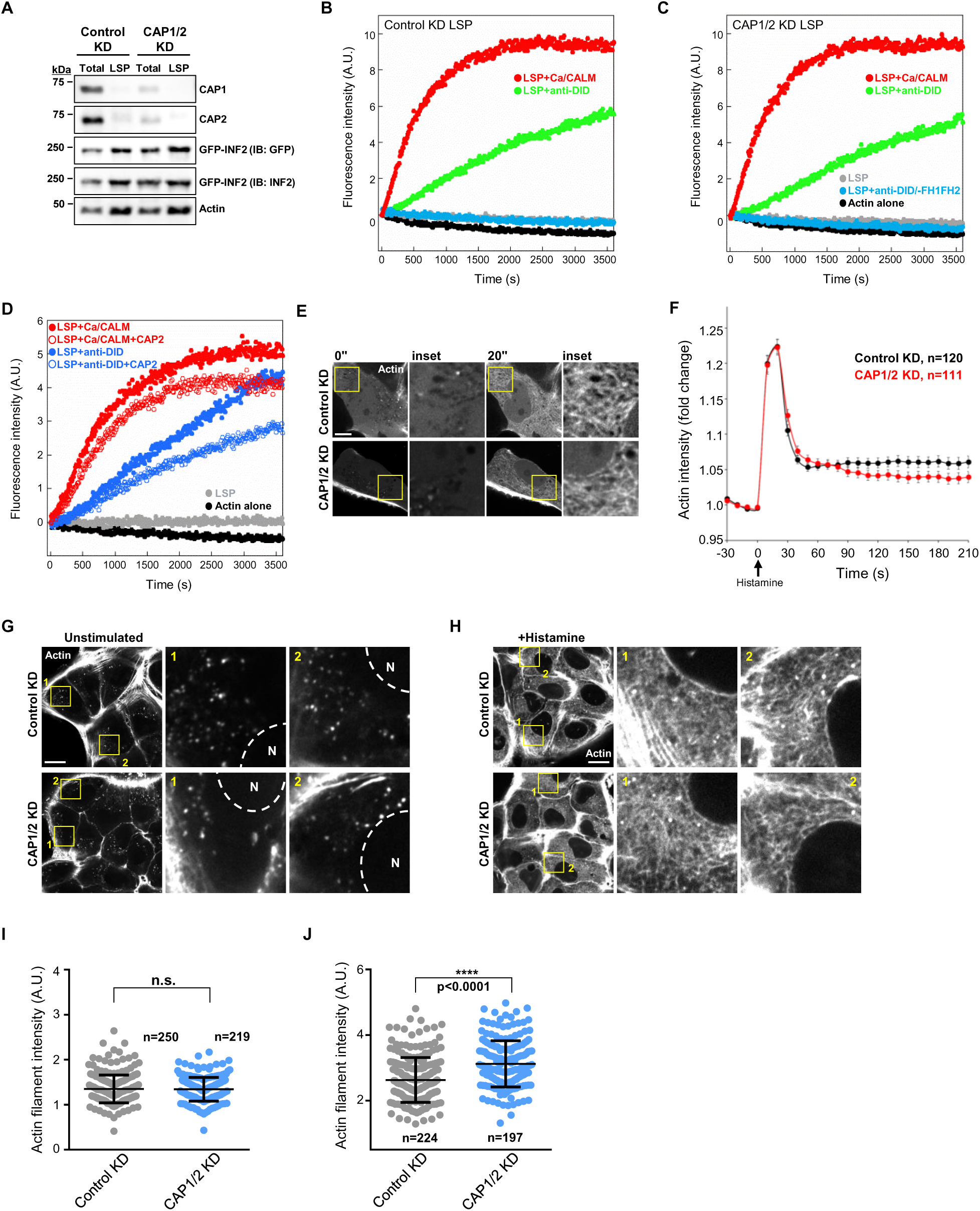
CAP proteins do not influence INF2 regulation. **(A)** Western blots of total cell lysate and LSP from U2-OS GFP-INF2-CAAX stable cells transfected with either control siRNA or CAP1/2 siRNAs. Equal protein (5 μg) was loaded in each lane and blotted with the indicated antibodies. **(B, C)** Pyrene-actin polymerization assays containing 2 μM actin (5% pyrene-labeled), 50 nM capping protein, 6 μM profilin, and 10 μg of LSP from U2-OS GFP-INF2-CAAX cells transfected with either control siRNA (B) or CAP1/2 siRNAs (C), with the indicated components (330 nM anti-DID antibody, 209 nM anti-FH1FH2 antibody, 1 μM calcium with 100 nM CALM). Experiments were performed four times with similar results, and the data shown are a representative experiment. **(D)** Pyrene-actin polymerization assay performed as in (B) and (C), without siRNA but including 1 μM recombinant CAP2 protein where indicated. Experiments were performed twice with similar results. Data shown are a representative experiment. **(E)** Time-lapse montage of U2-OS cells transfected with either control siRNA or CAP1/2 siRNAs, further transfected with mApple-F-tractin, and stimulated with 100 μM histamine at time 0. Insets show magnified views of the boxed regions. Scale bar: 10 μm; boxed region: 12 × 12 μm. **(F)** Time-course plot showing changes in cytoplasmic actin intensity (mApple-F-tractin) following histamine addition in U2-OS control KD cells and CAP1/2 KD cells. Y-axis values represent fluorescence intensity at each time point normalized to time 0 (F/F₀). Error bars represent the standard error of the mean (SEM), of the stated number of cells analyzed. **(G, H)** U2-OS cells transfected with either control siRNA or CAP1/2 siRNA were fixed and stained with TRITC-phalloidin without stimulation (G), or with 100 μM histamine treatment for 30 seconds (H). Insets show magnified views of the boxed regions, as indicated by the corresponding numbers. *N*=nucleus. Scale bar: 20 μm; boxed region: 20 × 20 μm. **(I, J)** Quantification of actin filament intensity in the perinuclear region without stimulation (H) or with 100 μM histamine treatment for 30 seconds (I), normalized to nuclear intensity. Each point represents one region of interest (ROI) per cell. *P*-value was calculated from an unpaired *t-*test. *n.s.*=not significant. error bars represent the standard error of the mean (SEM). Experiments were performed in triplicate.

We also tested the effect of CAP1/2 KD on CIA in intact U2OS cells. Upon histamine-stimulation, CAP1/2 KD cells display a similar time course of actin polymerization and depolymerization as control cells (**Fig. 5, E and F**). Additionally, we used TRITC-phalloidin staining of fixed cells as a means of examining cytoplasmic actin polymerization, which in our hands is more sensitive than the live-cell probes. No constitutive actin polymerization is detectable in CAP1/2 KD cells in the absence of histamine (**Fig. 5, G and I**). If CAP proteins were playing a role in suppressing unstimulated INF2 activity, their depletion should cause increased constitutive INF2 activity. Following histamine treatment, actin polymerization is slightly elevated in CAP1/2 KD cells compared to that observed in control KD cells (**Fig. 5, H and J)**.

To test whether this effect is cell-type specific, we also examined the impact of CAP1/2 KD on CIA in HeLa cells. Consistent with the findings in U2OS cells, CAP1/2 KD HeLa cells exhibit a similar time course of actin polymerization to control KD cells **(Fig. S3, A-C)**, and show no evidence of constitutive CIA in the absence of calcium stimulation, as assessed by fixed-cell imaging **(Fig. S3, D and F)**. Peak histamine-stimulated actin polymerization in CAP1/2 KD HeLa is elevated over control KD **(Fig. S3C)**. This modest increase in CIA intensity may reflect CAP’s known role in actin turnover (Kotila et al., 2018; Goode et al., 2023). Supporting this possibility, addition of CAP2 in the LSP assay results in slight inhibition of activated actin polymerization (**Fig. 5 D**).

Collectively, these results suggest that CAP1/2 proteins are not essential for INF2 regulation, in contrast with our previous findings (A et al., 2019, 2020). We next sought to answer how INF2 is tightly regulated in cells, while purified INF2 is constitutively active for actin polymerization.

### Effect of actin monomer binding proteins on INF2 activity

In previous studies, we showed that the DAD region of INF2 binds an actin monomer (Chhabra and Higgs, 2006), and that the actin/DAD interaction competes with the DID/DAD interaction (Ramabhadran et al., 2013). Addition of the actin monomer binding protein profilin, however, decreases INF2 activity in a DID-dependent manner, suggesting that actin monomer binding proteins are important for INF2 regulation (Ramabhadran et al., 2013). In mammalian cells, two abundant actin monomer binding proteins are profilin and thymosin (Pollard et al., 2000). We examined the relative abilities of these proteins to mediate INF2 regulation.

To test the effect of thymosin on INF2 regulation, we used pyrene-actin polymerization assays containing either full-length INF2 (INF2-FL) or the INF2-FFC construct, which lacks the DID (**Fig. 2 A**). The INF2 concentration used in these assays (10 nM) is in the same order of magnitude as the concentration we had measured in U2-OS cytoplasm (50 nM, A. et al., 2020). Titration of thymosin results in inhibition of INF2-FL at lower thymosin concentrations than for INF2-FFC (**Fig. 6, A and B, Fig. S4 A**). In both cases, there was a sigmoidal nature to the inhibition curves, and fitting to the Hill equation gave IC50s of 1.4 and 3.2 μM, and Hill coefficients of −0.53 and −0.65, respectively (**Fig. 6 B**). We next performed thymosin titration in the presence of 2 μM profilin, a concentration which only partially inhibits INF2-FL. Under this condition, the IC50 of thymosin drops to 0.6 μM and is non-sigmoidal (**Fig. 6, C and D**) while the IC50 for INF2-FFC is 1.9 μM with a Hill coefficient of −0.43 (**Fig. 6 D, Fig. S4 B**). These results suggest that, similar to profilin, thymosin-mediated inhibition of full-length INF2 occurs by competition with the DAD for actin monomer binding, allowing DAD to interact with DID.

**Figure 6.**
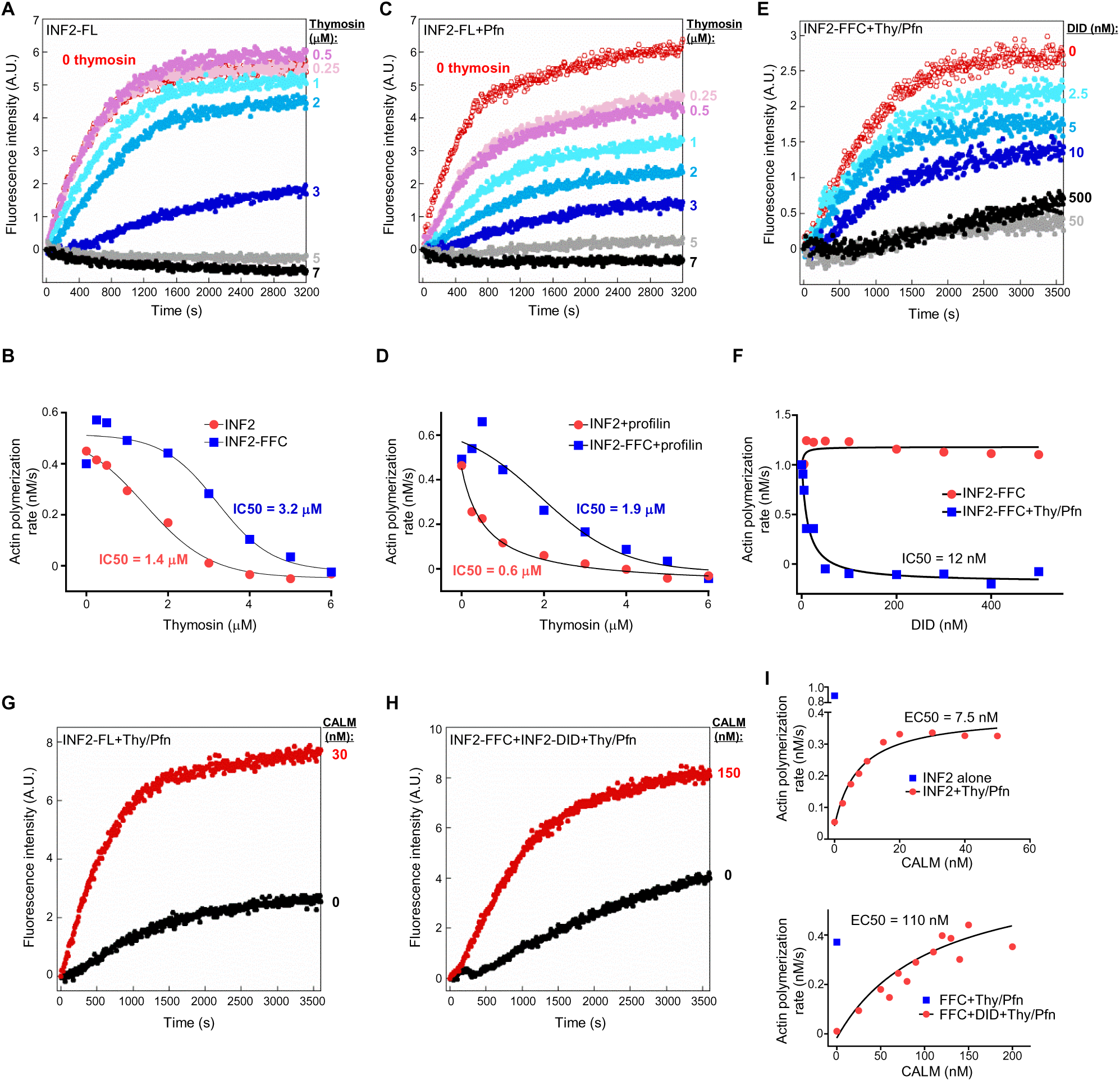
Actin monomer binding proteins allow reversible INF2 regulation. (A, **B)** Pyrene-actin polymerization assay containing 2 μM actin (5% pyrene-labeled), 50 nM capping protein, and 10 nM INF2 full-length (FL) protein, with the indicated concentrations of thymosin (A). Actin polymerization rates were calculated from the linear phase (0-250 seconds) of the pyrene-actin assay (B). Experiments were performed twice with similar results, and the data shown are a representative experiment. **(C, D)** Same as (A) and (B), but performed in the presence of 2 μM profilin. **(E, F)** Same as in (A), but performed in the presence of 10 nM INF2-FFC, 2.9 μM thymosin, 6 μM profilin and the indicated concentrations of INF2-DID (DID). Experiments were performed in triplicate with similar results, and the data shown are a representative experiment. **(G, H)** Pyrene-actin polymerization assay containing 2 μM actin (5% pyrene-labeled), 50 nM capping protein, 2.9 μM thymosin, 6 μM profilin, 1 μM calcium and either 10 nM INF2 full-length (G) or 10 nM INF2-FFC with 300 nM INF2-DID (H), in the presence of the indicated concentrations of CALM. **(I)** Calculations of actin polymerization rates for CALM concentration curves from experiments with INF2-FL (top) or INF2-FFC + INF2-DID (bottom).

We asked whether addition of INF2-DID would enable INF2-FFC inhibition in the presence of thymosin and profilin. In the absence of thymosin/profilin, a 50-fold excess of INF2-DID has minimal effect on INF2-FFC activity (**Fig. S4 C**), consistent with our past results (Ramabhadran et al., 2013). In contrast, low concentrations INF2-DID efficiently inhibit INF2-FFC in the presence of thymosin/profilin (**Fig. 6 E**), with an IC50 of 12 nM (**Fig. 6 F**).

We next tested the ability of calcium/CALM to activate INF2 in the presence of thymosin/profilin. For full-length INF2, calcium/CALM activates actin polymerization with an EC50 of 7.5nM (**Fig. 6 G and I**). For INF2-FFC in the presence of 300 nM INF2-DID, calcium/CALM activates actin polymerization with an EC50 of 110nM (**Fig. 6, H and I**). In the presence of lower INF2-DID concentration (50 nM), the CALM EC50 decreases to 10.2 nM (**Fig. S4 D**).

We also assessed the relative effects of thymosin and profilin in our cell-free system, using LSP and 50 nM capping protein. In the absence of thymosin or profilin, constitutive actin polymerization occurs (**Fig. S4 E**), as expected due to the lack of factors inhibiting nucleation or pointed end elongation. Titration of thymosin alone inhibits actin polymerization with an IC50 of 0.95 μM (**Fig. S4, E and G**), Addition of 0.25 μM profilin enhances the effect of thymosin, reducing its IC50 to 0.6 μM (**Fig. S4, F and G**). This inhibition of actin polymerization is reversed upon addition of either anti-DID antibody (**Fig. S4 H**) or calcium/CALM (**Fig. S4 I**).

Next, we tested whether a more physiological ratio of actin:capping protein:thymosin:profilin could recapitulate INF2 inhibition and activation in a cell-free system. To determine these ratios, we used published transcript levels from U20S cells (see Materials and Methods), calculating an actin:capping protein:thymosin:profilin of 1:0.034:1.2:0.27. In LSP assays containing 2 μM actin, 68 nM capping protein, 2.4 μM thymosin, and 0.54 μM profilin, actin polymerization is largely suppressed, but is potently activated by either anti-DID antibody or calcium/CALM **(Fig. S5 A)**. Increasing the levels of capping protein/thymosin/profilin by 1.5- or 2-fold relative to actin (while keeping the ratios of these three proteins the same) causes full inhibition of actin polymerization by LSP alone, which is again reversed by anti-DID or calcium/CALM treatment **(Fig. S5 B and C)**.

Taken together, these results suggest that actin monomer-binding proteins such as thymosin and profilin play a crucial role in controlling INF2 function. The interaction between INF2 DAD and DID regions is modulated by free actin monomer level, with monomer binding proteins reducing the concentration of free actin monomer to allow INF2 regulation. The experiment described in the next section test this hypothesis in cells.

### Actin monomer binding to the INF2 DAD is important for INF2 activation

To test the role of actin monomer binding by INF2-DAD in cellular INF2 regulation, we produced an INF2 construct possessing a DAD with reduced actin monomer binding affinity, by swapping INF2’s DAD with the DAD from the formin mDia1. In past studies, we have shown that mDia1-DAD binds to the INF2 DID with an affinity similar to that of INF2 DAD (Sun et al., 2011). However, mDia1 DAD exhibits a low affinity for actin monomers, compared to the INF2 DAD, which binds actin with nanomolar affinity (Chhabra and Higgs, 2006; Heimsath and Higgs, 2012; Gould et al., 2011). Using polarization assays, we confirm that the mDia1 DAD has significantly lower affinity for actin monomers compared to the INF2 C-terminal region, not reaching saturation at 20 μM actin, while maintaining similar affinity for INF2-DID (**Fig. 7 A**).

**Figure 7.**
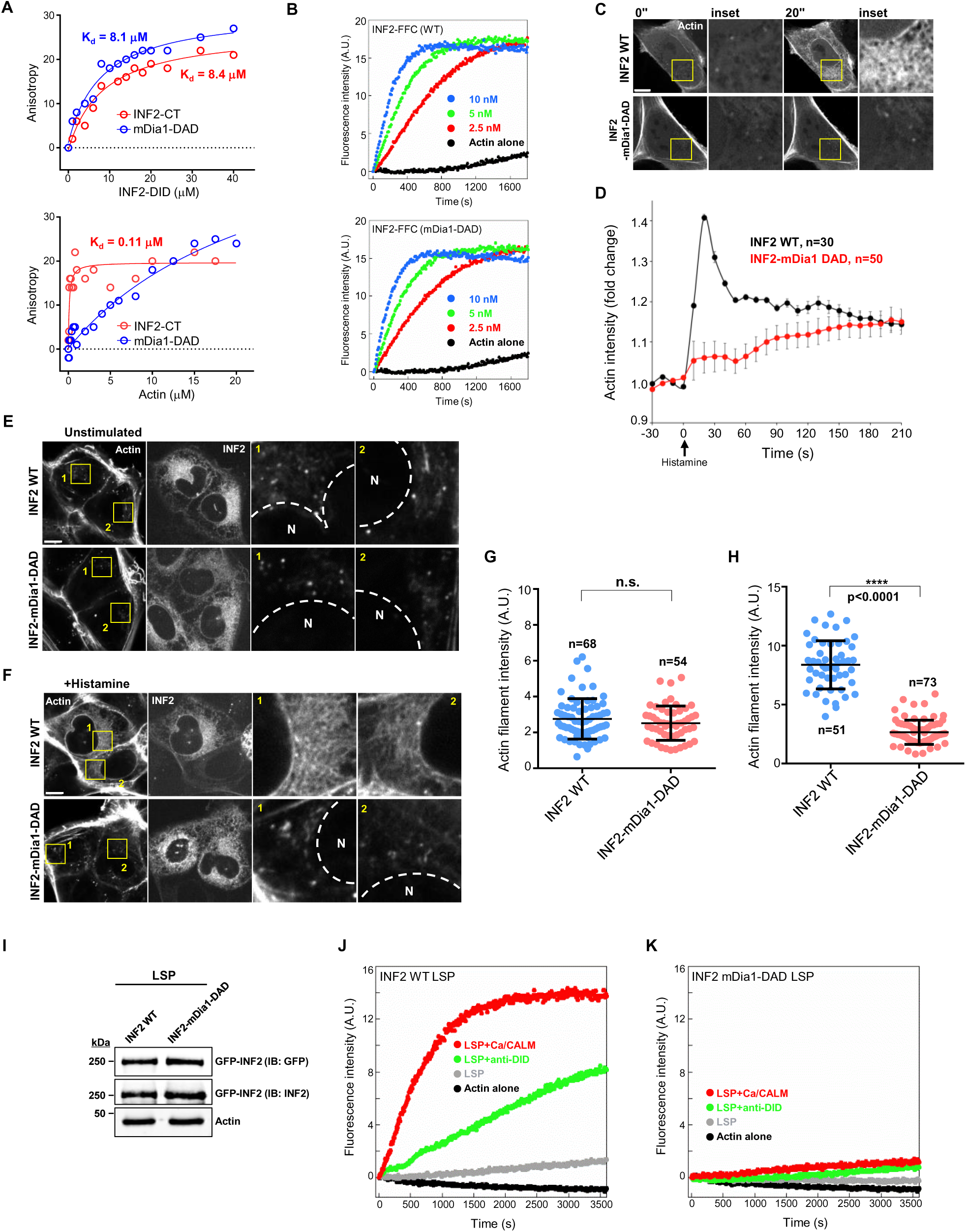
Replacing the INF2 DAD with the mDia1 DAD compromises INF2 activation. **(A)** Fluorescence anisotropy assay to measure actin monomer or INF2 DID binding by the INF2 C-terminal region or mDia1 DAD peptide. FITC-labeled INF2 C-terminal or TAMRA-labeled mDia1-DAD peptide (50 nM) were incubated with varying concentrations of LatA-stabilized actin or INF2 DID. **(B)** Pyrene-actin assays containing 2 μM actin monomers (5% pyrene labeled) and 2.5, 5, or 10 nM of the FFC construct of WT INF2 or the INF2-mDia1 DAD chimera. **(C)** Images from live-cell experiments in which U2-OS INF2 KO cells were transfected with mApple-F-tractin and either GFP-INF2 CAAX WT or GFP-INF2 CAAX-mDia1-DAD chimera. Cells stimulated with 100 μM histamine at time 0. Insets show magnified views of the boxed regions. Scale bar: 10 μm; boxed region: 12 × 12 μm. **(D)** Time-course plot showing changes in cytoplasmic actin intensity (mApple-F-tractin) following histamine addition in U2-OS INF2 KO cells expressing either GFP-INF2 CAAX-WT or GFP-INF2 CAAX-mDia1-DAD. Y-axis values represent fluorescence intensity at each time point normalized to time 0 (F/F₀). Error bars represent the standard error of the mean (SEM), of the stated number of cells analyzed. **(E, F)** U2-OS INF2 KO cells transfected with either GFP-INF2 CAAX-WT or GFP-INF2 CAAX-mDia1 DAD were fixed and stained with TRITC-phalloidin without stimulation (D), or with 100 μM histamine treatment for 30 seconds (E). Insets show magnified views of the boxed regions, as indicated by the corresponding numbers. *N*=nucleus. Scale bar: 10 μm; boxed region: 15 × 15 μm. **(G, H)** Quantification of actin filament intensity in the perinuclear region without stimulation (F) or with 100 μM histamine treatment for 30 seconds (G), normalized to nuclear intensity. Each point represents one region of interest (ROI) per cell. *P*-value was calculated from an unpaired *t-*test. *n.s.*=not significant. error bars represent the standard error of the mean (SEM). **(I)** Western blot of LSP from U2-OS INF2 KO cells transiently expressing either GFP-INF2-CAAX WT or GFP-INF2 CAAX-mDia1-DAD cells. Equal protein (5 μg) was loaded in each lane and blot was probed with the indicated antibodies. **(J, K)** LSP pyrene-actin polymerization assays containing 2 μM actin (5% pyrene-labeled), 50 nM capping protein, 6 μM profilin, and 10 μg of LSP from U2-OS INF2 KO cells transiently expressing GFP-INF2-CAAX wild-type (J) or GFP-INF2 CAAX-mDia1 DAD (K), with the indicated components (330 nM anti-DID antibody or 1 μM calcium with 100 nM CALM). Experiments were performed in triplicate with similar results, and the data shown are a representative experiment.

We then produced a chimeric construct, in which we replace INF2’s DAD with that of mDia1. In pyrene-actin assays, the FFC construct of this chimera displays a similar ability to stimulate actin polymerization as WT INF2-FFC (**Fig. 7B**). In cells, however, the full-length INF2–mDia1-DAD chimera displays greatly reduced histamine-activated actin polymerization compared to WT INF2 (**Fig. 7C, D**). Experiments using fixed-cell staining with TRITC-phalloidin confirm this result, showing that the INF2–mDia1-DAD chimera displays no apparent cytoplasmic actin polymerization either in the absence of stimulation or after 30 sec histamine stimulation (**Fig. 7 E-H)**. In the cell-free assay, LSP prepared from INF2-mDia1-DAD expressing cells exhibits low actin polymerization activity in the presence of anti-DID or calcium/CALM (**Fig. 7 I-K)**.

These findings suggest that actin monomer binding to the INF2 DAD region is an important step in cellular INF2 activation. Another possibility is that severing by INF2 is an important facet of CIA, as addressed in the Discussion.

### Importance of the INF2 N-terminal region to INF2 regulation

Formins of the Dia and FMNL classes possess clear Rho GTPase binding domains (GBD) N-terminal to the DID (Rose et al., 2005; Kühn et al., 2015). In contrast, INF2 contains a short (30 amino acid) sequence devoid of a GBD. However, a recent study showed that the INF2 N-terminus (NT) is a calmodulin binding site (Labat-de-Hoz et al., 2022). Using a fluorescently labeled INF2-NT peptide (FITC-INF2-NT) in fluorescence polarization assays, we confirm that INF2-NT directly binds CALM with a K_d_^app^ of ∼ 4 μM (**Fig. 8 A**), which is in the range previously determined by surface plasmon resonance (Labat-de-Hoz et al., 2022).

**Figure 8.**
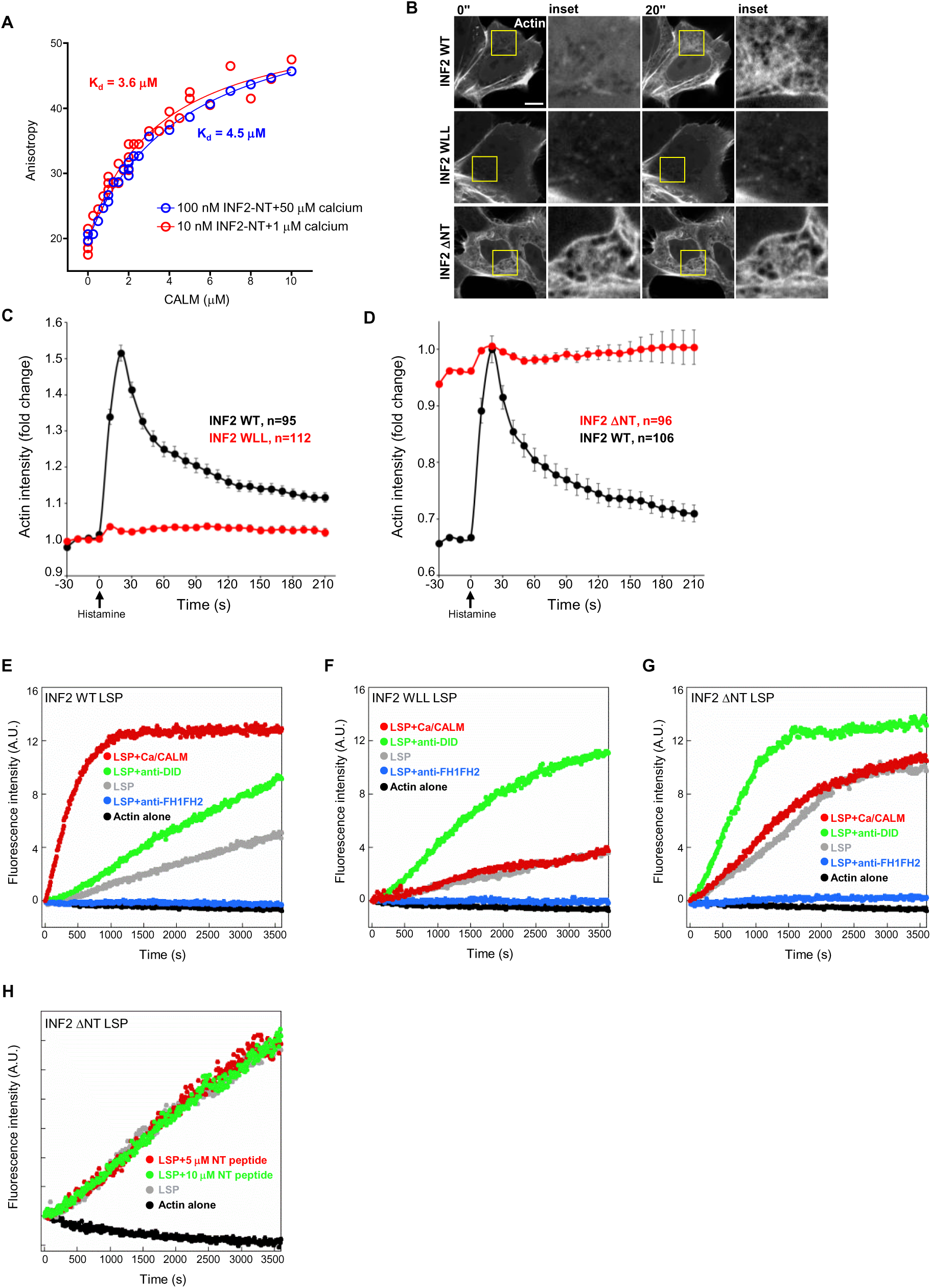
Deletion of the INF2 N-terminal region causes constitutive activity. **(A)** Fluorescence anisotropy assay to measure CALM binding by the INF2 N-term peptide (1-29 aa). FITC-labeled INF2 N-term was incubated with varying concentrations of CALM. Two conditions were used: Condition 1 was 100 nM FITC-INF2-NT and 50 μM free calcium (blue); Condition 2 was 10 nM FITC-INF2-NT and 1 μM free calcium (red). The K_d_^app^ were 4.5 and 3.6 μM, respectively. **(B)** Images from a time-lapse movie of U2-OS INF2 KO cells transiently expressing mApple-F-tractin along with either GFP-INF2-CAAX wild-type, WLL or ΔNT mutant. Cells were stimulated with 100 μM histamine at time 0. Insets show magnified views of the boxed regions. Scale bar: 10 μm; boxed region: 12 × 12 μm. **(C, D)** Time-course plot showing changes in cytoplasmic actin (mApple-F-tractin) following histamine addition in GFP-INF2-CAAX wild-type cells, WLL mutant (C) or ΔNT mutant (D). Y-axis values represent fluorescence intensity at each time point normalized to time 0 (F/F₀). Error bars represent the standard error of the mean (SEM) for indicated number of cells. **(E-G)** Cell-free pyrene-actin polymerization assays containing 2 μM actin (5% pyrene-labeled), 50 nM capping protein, 6 μM profilin, and 10 μg of LSP from U2-OS INF2 KO cells transiently expressing GFP-INF2-CAAX wild-type (E), WLL mutant (F) or ΔNT mutant (G), with the indicated components (330 nM anti-DID antibody, 209 nM anti-FH1FH2 antibody, or 1 μM calcium with 100 nM CALM). Experiments were performed in triplicate with similar results, and the data shown are a representative experiment. **(H, I)** Cell-free pyrene-actin polymerization assays containing 2 μM actin (5% pyrene-labeled), 50 nM capping protein, 6 μM profilin, and 10 μg of LSP from U2-OS INF2 KO cells transiently expressing GFP-INF2-CAAX wild-type (H), or ΔNT mutant (I), with the indicated components (5 μM or 10 μM INF2 N-terminal peptide).

To evaluate the importance of the calmodulin-binding site for INF2 activation, we tested several INF2-CAAX mutants for unstimulated and histamine-stimulated actin polymerization when transiently expressed in INF2-KO U2OS cells. All mutants display similar expression levels and ER localization to WT INF2 (**Fig. S6**). As found in (Labat-de-Hoz et al., 2022), a triple mutation of W11, L14, and L18 to alanines (INF2-WLL) renders INF2 inactive in both the unstimulated and histamine-stimulated states (**Fig. 8 B and C**). Similar results occur upon mutation of four basic residues in this region (R9, K10, K15, K17), whereas single mutants of R9, A13 or E16 do not alter basal or histamine-stimulated INF2 activity (**Fig. S7 A**). While a bacterially-expressed INF2 construct containing the N-terminus (1-420, **Fig. 2 A**) demonstrates competition with FITC-INF2-NT for CALM binding, the corresponding WLL mutant does not compete (**Fig. S7 B**), demonstrating effective ablation of CALM binding by the WLL mutation.

We also tested an INF2 construct lacking the N-terminal 30 amino acids (INF2 ΔNT). Similar to INF2-WLL, INF2 ΔNT does not compete with FITC-INF2-NT peptide for CALM binding (**Fig. S7 B**). Our expectation was that this construct would behave similarly to INF2-WLL in cells. Surprisingly INF2 ΔNT displays cytoplasmic actin filaments even in unstimulated cells, suggesting constitutive activation similar to the INF2-R218Q mutant (**Fig. 8, B and D**).

We used the cell-free assay to examine the WLL and ΔNT mutants further, comparing against INF2-WT which displays robust activation by calcium/CALM (**Fig. 8 E**). For INF2-WLL, actin polymerization is low with LSP alone and is activated by anti-DID, but not by calcium/CALM (**Fig. 8 F**). These results are consistent with the hypothesis that CALM binding to the N-terminus is required for INF2 activation. In contrast, LSP from INF2 ΔNT cells displays high constitutive actin polymerization in the absence of activator, with calcium/CALM not further activating (**Fig. 8 G**).

These results suggest that INF2’s N-terminus serves as more than simply a CALM binding site, and may act in the autoinhibitory interaction itself. However, addition of INF2 N-terminal peptide to the LSP does not inhibit constitutive actin polymerization LSP from INF2 ΔNT cells (**Fig. 8 H).** An alternative possibility is that deletion of the N-terminus compromises DID/DAD binding. For this reason, we used fluorescence polarization to test the affinities of INF2 constructs ending at amino acid 420 (**Fig. 2 A**) for INF2’s DAD-containing C-terminal region. All three constructs (WT, WLL and ΔNT) bind INF2-Cterm with similar affinity (K_d_^app^ between 2.6 and 4.1 µM) (**Fig. S7 C**).

The INF2 N-terminal region is highly conserved, with the first 30 amino acids being identical in placental mammals (**Fig. S7 D**). A previous NMR-based study (Labat-de-Hoz et al., 2022) reported that the N-terminal region is composed of two helices connected by a short linker (amino acids 20– 24). We designed a mutant (INF2-LM) that replaces this linker sequence with the generic linker GSGSG. We also created a second mutant replacing amino acids 1-5 with the same sequence (INF2-NM). We then tested the effects of these mutations on unstimulated and histamine-stimulated actin polymerization when expressed as the GFP-INF2-CAAX construct in INF2-KO U2OS cells. While INF2-NM displays similar activity to INF2-WT, the INF2-LM mutant is less tightly regulated in three respects. First, INF2-LM cells display constitutive actin polymerization in the peri-nuclear region (**Fig. 9 A**). The degree of this constitutive activity is mild compared to the R218Q or ΔNT mutants, as evidenced by a minor increase in actin intensity pre-stimulation, and the ability of histamine to stimulate a further increase (**Fig. 9 B**). We confirmed this constitutive actin by fixed-cell staining with TRITC-phalloidin, and found a significant increase in unstimulated peri-nuclear actin intensity (**Fig. 9, C and D**). Second, INF2-LM expressing cells display delayed actin filament disassembly after histamine stimulation (**Fig. 9, A and B**). Third, in the cell-free assay the INF2-LM mutant exhibits greater actin polymerization activity in the absence of any activators, with calcium/CALM further enhancing actin polymerization (**Fig. 9, E and F**). Interestingly, the INF2-LM 1-420 construct can still compete with fluorescently-labeled INF2-NT peptide for CALM binding (**Fig. S7 B**) and binds to INF2-Cterm with similar affinity to WT (**Fig. S7 C**). Taken together, these findings suggest that the linker region plays a role in maintaining INF2 inhibition.

**Figure 9.**
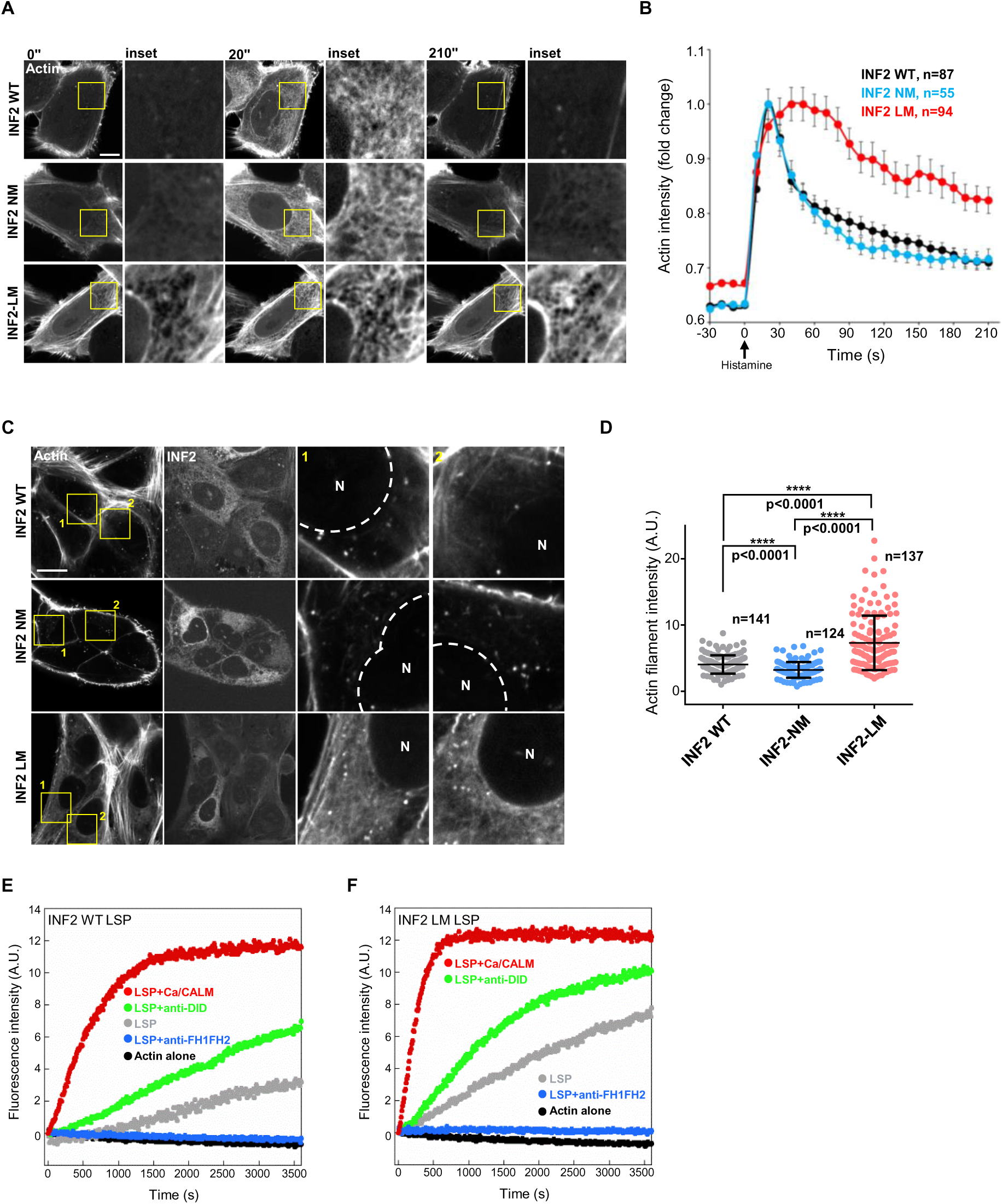
The N-terminal linker region of INF2 is required for ebicient regulation. **(A)** Time-lapse montage of U2-OS INF2 KO cells transiently expressing mApple-F-tractin along with either GFP-INF2-CAAX wild-type, NM or LM mutant. Cells were stimulated with 100 μM histamine at time 0. Insets show magnified views of the boxed regions. Scale bar: 10 μm; boxed region: 12 × 12 μm. **(B)** Time-course plots showing changes in cytoplasmic actin filaments (mApple-F-tractin) following histamine addition in GFP-INF2-CAAX wild-type, NM or LM mutant expressing cells. Y-axis values represent fluorescence intensity at each time point, normalized to the maximum fluorescence intensity observed. Error bars represent the standard error of the mean (SEM) of the indicated cell numbers. **(C)** U2-OS INF2 KO cells transiently expressing GFP-INF2-CAAX wild-type, NM or LM mutant were fixed and stained with TRITC-phalloidin without stimulation. Insets show magnified views of the boxed regions. *N*=nucleus. Scale bar: 20 µm; boxed region: 20 × 20 µm. **(D)** Quantification of actin filament intensity in the perinuclear region, normalized to nuclear intensity. Each point represents one region of interest (ROI) per cell. *P*-value calculated from an unpaired *t-*test. Error bars represent the standard error of the mean (SEM). **(E,F)** Cell-free pyrene-actin polymerization assays containing 2 μM actin (5% pyrene-labeled), 50 nM capping protein, 6 μM profilin, and 10 μg of LSP from U2-OS INF2 KO cells transiently expressing either GFP-INF2-CAAX wild-type (E), or LM mutant (F), with the indicated components (330 nM anti-DID antibody, 209 nM anti-FH1FH2 antibody, or 1 μM calcium with 100 nM CALM). Experiments were performed in triplicate with similar results, and the data shown are a representative experiment.

## Discussion

In this work, we develop a facile cell-free system (called the LSP assay) to assess INF2 activity, involving the enrichment of ER membrane (containing bound INF2-CAAX). The rapidity of this preparation (requiring <30 min) allows for a broad range of assays. The removal of cytosolic components (actin monomers, actin binding proteins, nucleotides, inorganic ions) allows testing of specific factors by add-back. We use the system to test INF2 activation, and show that either a DID-directed antibody or calcium-bound calmodulin (CALM) are potent activators. The specificity of these stimuli for INF2 are suggested by two results: 1) INF2-KO cells display no response to either anti-DID or calcium-CALM, and 2) an FH1FH2-directed antibody inhibits this activation. In view of the fact that several other formins are enriched in this extract (mDia2, FMNL1-3), their regulation might also be studied in this system. It should be noted that we use an actin monomer concentration (2 μM) that is far below the cytoplasmic level (estimated at 50 μM, see Results). Further honing of this system will require increasing this concentration, in the context of increased actin monomer binding proteins to prevent spontaneous polymerization.

Assays using this cell-free system, as well as assays with purified proteins, support the following model of INF2 regulation (**Fig. 10**). In unstimulated cells, INF2 is maintained in an inactive state by DID/DAD interaction. The integrity of this autoinhibitory interaction requires the presence of actin monomer-binding proteins (such as profilin or thymosin) to compete with DAD for actin monomer binding. INF2 activation occurs through disruption of the DID/DAD interaction by calcium-CALM interaction with the N-terminal region of INF2. We discuss key aspects of the model, its relationship to previous studies, and puzzling open questions below.

**Figure 10.**
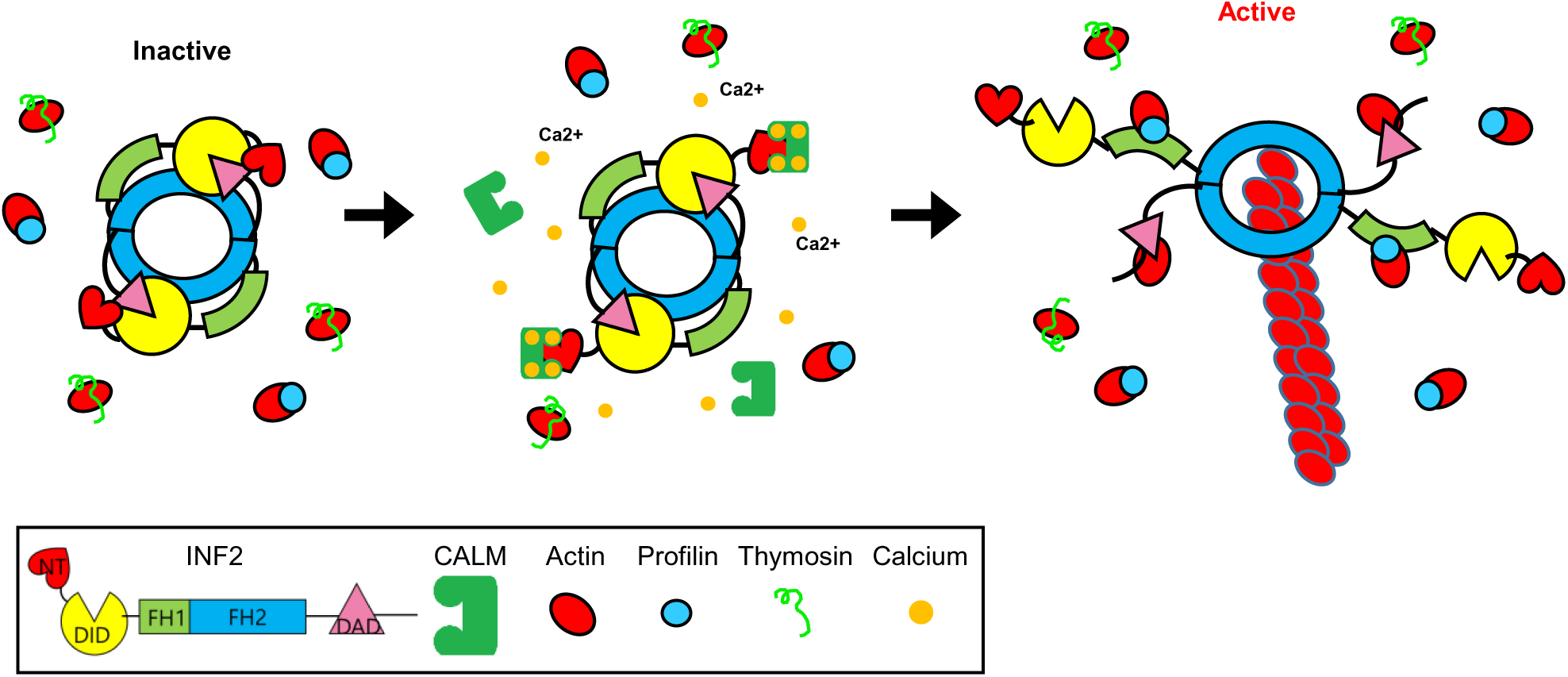
Model of INF2 regulation. INF2 (a dimer) is autoinhibited by the DID/DAD interaction, which also requires actin monomers to be bound by monomer-binding proteins like profilin or thymosin. Increased calcium enables calcium-calmodulin (CALM) binding to INF2’s N-terminus, which disrupts the DID/DAD interaction. Actin monomer binding by the DAD keeps INF2 in the activated state.

The model is built on our results as well as previous results showing that: 1) INF2 is activated by increased calcium in cells (Shao et al., 2015; Wales et al., 2016; Chakrabarti et al., 2018), 2) INF2 binds calcium-CALM (Wales et al., 2016), 3) calcium-CALM binding is necessary for INF2 activation in cells (Labat-de-Hoz et al., 2022), and actin monomers compete with DID for DAD binding (Ramabhadran et al., 2013). The novelty of our study is that we put these results together in the context of both purified proteins and a cell-free assay.

The activation of INF2 by calcium-CALM is rapid, transient, and robust. In cells, the increase and decrease of CIA closely parallels the rise and fall of cytoplasmic calcium, with actin dynamics lagging behind calcium dynamics by less than 5 sec (Chakrabarti et al., 2018). We show here that INF2 mobilizes a significant population of the cell’s actin in a narrow time window. In response to ionomycin, ∼31% of cellular actin is converted from monomer to polymer within 30 sec, while histamine causes ∼21% conversion within 10 sec. INF2 KO or KD abolishes CIA (Wales et al., 2016; Chakrabarti et al., 2018; Fung et al., 2019). It is possible that other actin polymerization mechanisms contribute to this impressive total of polymerized actin. However, we have previously shown that Arp2/3 complex inhibition does not affect the timing or extent of CIA (Fung et al., 2019), so another polymerization factor would have to be responsible. We think it more likely that our calculation is an underestimate of CIA-polymerized actin, because a previous study showed that calcium also triggers rapid actin rapid disassembly of cortical actin filaments on the same time scale as CIA (Wales et al., 2016).

The mechanism by which INF2 mediates cellular actin polymerization is not clear. While being a potent nucleation factor, INF2 also is a potent actin filament severing protein (Chhabra and Higgs, 2006), requiring the actin to release its bound phosphate product of ATP hydrolysis before severing (Gurel et al., 2014a). In addition, INF2 allows actin filament elongation even in the presence of capping protein (Gurel et al., 2015), similar to other formin proteins. The percentage of cellular actin polymerization that is due to new filament nucleation, severing existing filaments, or mediating elongation of existing filaments is unknown. Another question is whether competition between INF2-DAD and other binding proteins (profilin, thymosin) for actin monomers would affect DAD’s participation severing or barbed end elongation. A recent study elegantly shows how INF2’s DAD participates in the severing process (Palmer et al., 2024). Removal of an actin monomer from DAD might in fact enhance severing. Given this ambiguity about the cellular relevance of severing to CIA, our INF2 chimera experiment (Fig. 7) could be explained by a severing defect. Unfortunately, our efforts to design INF2 mutants defective in severing but still maintaining auto-inhibitory regulation have not been successful to date.

In both our cell-free assay and assays using purified proteins, nanomolar CALM and sub-micromolar calcium are sufficient for INF2 activation, strongly suggest that calcium-CALM is the dominant activator of INF2. It is interesting that the dissociation constant for direct binding of INF2’s N-terminus to calcium/CALM (∼4 μM) is significantly higher than the concentration needed to activate INF2-mediated polymerization in either the LSP assay (EC50 9 nM) or with purified proteins (EC50 7.5 or 110 nM, depending on the conditions). This difference might be due to a difference in affinity of calcium/CALM for the minimal N-terminal peptide versus the full-length protein. Alternately, the difference might be a reflection of the difference between an equilibrium-based binding assay and the polymerization assay (in which, once a filament is nucleated, it continues to contribute to the signal whether or not INF2 subsequently releases from CALM).

In addition to activation by calmodulin, INF2 activity is sensitive to free actin monomer concentration. This is an intriguing case of an actin assembly factor integrating monomer availability into its activation mechanism. This mechanism might apply to other actin assembly factors, including other formins for which the DAD has affinity for monomers. INF2’s DAD binds actin monomer with high affinity, with K_d_ 40-120 nM (this publication, Chhabra & Higgs, 2006; Ramabhadran et al., 2013). The DAD of mDia1 has significantly lower affinity than INF2-DAD for actin monomers (this publication), but has been shown to contribute to actin dynamics (Gould et al., 2011). Members of the FMNL formin sub-family bind actin monomers at a site between the FH2 and the DAD, with a Kd of 0.9 μM (Heimsath and Higgs, 2012; Vaillant et al., 2008). The *Drosophila* DAAM formin also interacts with actin monomers, through a region C-terminal to the DAD (Vig et al., 2017), but this interaction is not competitive with profilin. Finally, the C-terminal region of the *Drosophila* Cappucino formin both interacts with actin monomers and is involved in regulation, although its sequence is not similar to DAD (Bor et al., 2012; Vizcarra et al., 2014). Whether the affinities of these interactions with actin are high enough to influence activation of these formins remains to be tested.

Two key questions to consider regarding the regulatory potential for actin monomers are: 1) what is the cytoplasmic concentration of free actin monomers (since binding by profilin, thymosin, or other monomer binding proteins is likely to compete with DAD), and 2) does this concentration fluctuate in cells? The fact that actin monomer level regulates transcription through MRTF proteins (Posern and Treisman, 2006) suggests that monomer levels do indeed fluctuate. A previous study showed that CIA occurs simultaneously with depolymerization of actin filaments near the plasma membrane (Wales et al., 2016), with the cortical depolymerization perhaps enhancing INF2 activation. An increase in actin monomers might, therefore, synergize with activation by calcium-CALM, making the CIA response more robust. In this respect, actin monomer-based regulation of INF2 might represent another facet to the concept of ‘competition’ between actin-based structures, whereby inhibition or activation of one actin-based structure can influence the dynamics of a second structure through monomer availability (Kadzik et al., 2020). It is possible that the transient nature of CIA is due to depletion of actin monomers, which then shifts the equilibrium from DAD/actin to DAD/DID interaction.

Our findings also emphasize the importance of actin monomer ‘buffers’ in the cytoplasm, and our knowledge gaps in several respects. First, while we treat profilin and thymosin as single entities here, there are multiple genes for both (four and five, respectively) in mammals. Transcriptomic data suggest that profilin 1 (PFN1, the profilin we use here) is predominant in U2-OS cells (1244 transcripts per million (TPM), versus 164, 0, and 1.11 respectively for PFN2, PFN3 and PFN4), but past studies have shown significant differences between formin interactions with specific profilins (Neidt et al., 2009). For thymosins, both thymosin beta-4 and beta-10 are highly expressed in U2OS cells (3237 and 2973 TPM, respectively), yet only thymosin beta-4 has been well characterized. Second, it is unclear if other actin binding proteins can contribute to monomer buffering. Third, more detailed information on relative cellular protein levels of actin, profilin, thymosin, and other monomer binding proteins are needed for a full understanding of monomer buffering.

Our current results argue strongly against our previously proposed model, in which cyclase-associated proteins (CAPs) mediated an inhibitory complex, along with acetylated actin (A et al., 2019, 2020). While the combination of CAP and actin inhibits INF2 in certain situations in the context of purified proteins, it does not play a role in our cell-free system. In addition, actin monomer buffering by profilin and/or thymosin is sufficient to allow INF2 autoinhibition with purified proteins. Interestingly, CAP1/2 KD also has no appreciable effect on the kinetics of CIA depolymerization after histamine stimulation in U2-OS cells, but does slow CIA depolymerization somewhat in HeLa cells. This result raises questions as to the factors controlling the rapid turnover of actin filaments during CIA, and whether these factors vary by cell type. For example, CAP proteins have been shown to counter-act INF2 activity during dendritic spine maturation (Schuldt et al., 2024). CAP is one of several proteins that has powerful effects on actin turnover, with others being cofilins, Aip1, coronins, and twinfilins (Goode et al., 2023). The combined effects of these proteins are often more potent than those of the individual proteins, so a combination of turnover factors is probably required. It is also possible that the potent severing activity of INF2 (Gurel et al., 2014a) might itself play a role in the depolymerization phase of CIA. It will be interesting to include a greater suite of actin turnover proteins in the LSP assay, to recapitulate the depolymerization phase.

Two results suggest that the N-terminal region of INF2 serves as more than just a calmodulin binding site. First, deletion of the N-terminus results in constitutively active INF2, while point mutants eliminating calmodulin interaction cause a permanently ‘off’ INF2 that is not activatable in cells. Second, substituting a region outside of the calmodulin binding site (amino acids 20-24, PQDSD) with a flexible linker (GSGSG) also causes constitutively active INF2. One possible explanation is that the N-terminal region interacts with another region of INF2 in the inhibited state. Indeed, one AlphaFold model of an INF2 construct lacking the DAD region (A0A087X118) predicts that the N-terminus can occupy a site on the DID that overlaps with the DAD-binding site (**Fig. S7 E-H**). Molecular dynamics simulations based on this model demonstrated stable binding of the N-terminus within this site over at least 100 ns of simulation at 310 K (see Materials and Methods for details), indicating that this predicted interaction is energetically favorable. While these models are speculative, they raise the possibility of additional N-terminal interactions that influence INF2 regulation. While we do not observe effects of the N-terminus on DID/DAD interaction by fluorescence anisotropy, a possibility is that the effect of the N-terminus depends on INF2 being dimeric, as in the full-length protein.

Overall, our results suggest an elegant and efficient mechanism for INF2 regulation, with actin monomer buffering being required to allow INF2 autoinhibition, and calcium-calmodulin causing rapid activation. INF2 activation results in a burst of actin polymerization that consumes 20-30% of total cellular actin within seconds. Considering the fact that mutations to over 40 residues in the DID region are linked to human disease (Labat-de-Hoz and Alonso, 2020), this regulation is clearly important for cellular homeostasis.

## Materials and methods

### Plasmids and siRNA oligonucleotides

The GFP-F-tractin and mApple-F-tractin plasmids were gift from A. Pasapera (NIH, Bethesda, MD). The ER-tagRFP (modified from the GFP-N1 backbone) was a gift from E. Snapp (Albert Einstein College of Medicine, New York, NY); and contains a prolactin signal sequence at the 5’ end of the fluorescent protein as well as a KDEL sequence at the 3’ end. The GFP-fusion constructs of human INF2-CAAX (UniProt Q27J81-1) and INF2-nonCAAX (UniProt Q27J81-2) constructs used in this study have been describe previously (Chakrabarti et al., 2018; A et al., 2019), and are in a modified eGFP-C1 vector (Clontech) containing a Strep-tag II (IBA Life Sciences) and an HRV3C cleavage site between the multiple cloning site and the start codon of INF2. INF2 mutations (R218Q; W11L14L18 to A; ΔNT; NM; LM; R9A, R9K10K15K17 to A; A13T; and E16A) were introduced into the GFP-INF2-CAAX construct using site-directed mutagenesis. For bacterial expression, human INF2 FH1-FH2-C (FFC, amino acids 469-1250) has been described previously (Ramabhadran et al., 2012). Human INF2 (amino acids 1–420) has been described previously (Sun et al., 2011). Mutants (W11L14L18 to A, ΔNT, and LM) were generated in the wild-type construct using site-directed mutagenesis. Human INF2-C term (amino acids 941-1249) was PCR amplified from GFP-INF2-CAAX and cloned into a pHis6-parallel1 plasmid (NovoPro Bioscience; V005547). His6-TEV-Hs CALM (residue 2-148) was obtained from addgene (159693) and has been described previously (Agamasu et al., 2019). The human thymosin β4 (Uniprot ID: P62328) open reading frame was made by gene synthesis and cloned into a pMW plasmid. The codon-optimized insert is 5’-ATGAGCGATAAACCGGATATGGCGGAAATTGAAAAATTTGATAAAAGCAAACTGAAAAAAACCGAAACCCAGGAAAAAAACCCGCTGCCGAGCAAAGAAACCATTGAACAGGAAAAACAGGCGGGCGAAAGCTGA-3’ (Integrated DNA Technologies). Oligonucleotides used for siRNA mediated gene silencing (Integrated DNA Technologies): CAP1 (exon 12): 5’-GTCAGTGCCAAATCTTCC-3’, CAP2 (3’ UTR): 5’-CTTTGAGAATCTAAGATG-3’.

### Cell culture

Human osteosarcoma U2-OS cells (ATCC HTB-96) were cultured in Dulbecco’s Modified Eagle Medium (DMEM; Corning, 10-013-CV) supplemented with 10% newborn calf serum (R&D Systems; S11250). Human cervical cancer HeLa cells (ATCC CCL-2) were cultured in DMEM supplemented with 10% fetal bovine serum (Sigma-Aldrich; F4135). INF2 knockout (KO) U2-OS cells were generated in-house using CRISPR/Cas9 technology, as previously described (Chakrabarti et al., 2018). A U2-OS INF2 KO cell line stably expressing GFP-INF2-CAAX was made by transfecting the GFP-INF2-CAAX plasmid described above into INF2-KO U2-OS cells, then selecting cells with G418 (Sigma-Aldrich; A1720). G418-resistant cells were then enriched for GFP expression by multiple rounds of flow sorting (Dartmouth Flow Cytometry Core) to establish a cell population in which >90% of cells were GFP-positive. Cells were periodically flow sorted thereafter to maintain enrichment. All cell lines were maintained at 37 °C in a humidified incubator with 5% CO₂. Cells were assayed every 6 months for mycoplasma using the MycoStrip kit (InvivoGen; rep-mys-50).

### Transfections

For plasmid transfections, cells were seeded at 5 × 10⁵ cells per 35-mm well approximately 18 hours before transfection. Transfections were performed in OPTI-MEM medium (Life Technologies; 31985062) using 2 µl Lipofectamine 2000 (Invitrogen; 11668) per well for 3.5 hours. Cells were then trypsinized and replated onto glass-bottom dishes (MatTek Corporation; P35G-1.5-14-C) at a density of ∼3.3 × 10⁵ cells per well. Imaging was performed ∼24 hours post-transfection in live-cell imaging medium (DMEM supplemented with 25 mM D-glucose, 4 mM L-glutamine, 25 mM HEPES, and 10% newborn calf serum). For all experiments, the following amounts of DNA were transfected per well (individually or combined for cotransfections): 400 ng for GFP-F-tractin, 400 ng for mApple-F-tractin, 400 ng for ER-tagRFP, and 200 ng for GFP-tagged INF2-CAAX constructs.

For siRNA transfections, 1 × 10⁵ cells were seeded per well in six-well plates. Each well received 2 µl RNAiMAX (Invitrogen; 13778) and 63 pg of siRNA. Transfections were repeated after 48 hours, and cells were analyzed 96 hours after the original transfection.

For live-cell imaging experiments following siRNA treatment, mApple-F-tractin was transfected at 72 hours post-siRNA treatment as described above. 3.3 × 10⁵ cells were replated onto 35-mm glass-bottom dishes and imaged at 96 hours post-transfection.

### Primary Antibodies

Anti-actin (mouse, mab1501R; Millipore) was used at 1:10,000. Anti-tubulin (mouse, T9026; Millipore) was used at 1:10,000. Anti-GFP (mouse, A11120; Thermo Fisher) was used at 1:1,000. Anti-calnexin (rabbit, Cell Signaling Technology; 2433) was used at 1:1,000. Anti-ATP synthase subunit β (mouse, A21351; Thermo Fisher Scientific) was used at 1:2,000. Anti-GM130 (rabbit, ab52649; Abcam) was used at 1:1,500. Anti-GAPDH (mouse, sc-365062; Santa Cruz Biotechnology) was used at 1:3,000. Anti-profilin1 (rabbit, P7624; Millipore) was used at 1:2,000. Anti-cofilin (rabbit, 5175; Cell Signaling Technology) was used at 1:1,000. Anti-FMNL1 (rabbit, HPA008129; Millipore) was used at 1:1,000. Anti-FMNL2 (rabbit, HPA005464; Millipore) was used at 1:1,500. Anti-FMNL3 (rabbit, NBP2-24724; Novus Biologicals) was used at 1:1,000. Anti-Arp2 (mouse, ab129018; Abcam) was used at 1:500. Anti-Arp3 (mouse, sc-48344; Santa Cruz Biotechnology) was used at 1:1,000. Anti-Myosin IIA (rabbit, 3403; Cell Signaling Technology) was used at 1:1,000. Anti-Myosin IIC (rabbit, 8189; Cell Signaling Technology) was used at 1:1,000. Anti-fascin (mouse, SC-21743; Santa Cruz Biotechnology) was used at 1:1,000. Anti-actinin 1 (mouse, 05-384; Millipore) was used at 1:1,000. Anti-actinin 4 (rabbit, 15145s; Cell Signaling Technology) was used at 1:1,000. Anti-calmodulin (rabbit, D1F7J; Cell Signaling Technology) was used at 1:1,000. Anti-INF2 was described previously (Ramabhadran et al., 2011), amino acid 469-1249, rabbit) and was used at 1:1,000. Polyclonal antibody against bacterially-expressed mouse mDia1 amino acids 1-549 (expression/purification described in (Li and Higgs, 2003) was raised in chicken (Cocalico Biologicals) and affinity purified by using the antigen coupled to Sulfolink (Pierce, Thermo Fisher Scientific, Rockford, IL). The rabbit polyclonal antibody against mouse mDia2 (also called DIAPH3) amino acids 1-520 was described in (Hager et al., 2012), and referred to as HNH3.1 in that publication. Rabbit polyclonal antibodies against CAP1 and CAP2 were described in (A et al., 2020). Rabbit polyclonal antibody against human INF2 FH1-FH2 region (amino acids 469-940) was previously described (Ramabhadran et al., 2011). Polyclonal antibody against mouse INF2 (Q0GNC1) N-terminal region (containing the DID) amino acids 1-424 was raised in rabbits (Cocalico Biologicals) and affinity purified by using the antigen coupled to Sulfolink (Pierce, Thermo Fisher Scientific, Rockford, IL).

### Western blotting

For analysis of whole-cell extract preparations, confluent cells in 35-mm dishes were lysed in ∼300 µl of 1× DB buffer (50 mM Tris-HCl, pH 6.8; 2 mM EDTA; 20% glycerol; 0.8% SDS; 0.02% bromophenol blue; 1 M NaCl; 4 M urea), followed by heating at 95 °C for 5 min. Genomic DNA was sheared by passing the lysate through a 27G needle. Proteins were resolved by standard SDS–PAGE and transferred onto PVDF membranes (Millipore). Membranes were blocked in TBS-T (20 mM Tris-HCl, pH 7.6; 136 mM NaCl; 0.1% Tween-20) containing 3% bovine serum albumin for 1 h at room temperature, then incubated overnight at 4 °C with primary antibodies. After washing with TBS-T, membranes were incubated with horseradish peroxidase (HRP)-conjugated secondary antibodies (goat anti-mouse, #1721011, Bio-Rad; goat anti-rabbit, #1706515, Bio-Rad; rabbit anti-chicken/turkey, #613120, Molecular Probes) for 1 h at room temperature. Detection was performed using ECL Prime Western Blotting Detection Reagent (45-002-40, Cytiva Amersham), and chemiluminescent signals were captured with an ECL Chemocam imaging system (SYNGENE G:BOX Chemi XRQ). For fluorescent Western blotting, membranes were incubated for 1 h at room temperature with IRDye-conjugated secondary antibodies: IRDye 680 goat anti-mouse (926-68070, LI-COR) or IRDye 800CW goat anti-rabbit (926-32211, LI-COR). Signals were detected using the LI-COR Odyssey CLx imaging system.

### Protein expression and purification

GFP-INF2-nonCAAX (plasmid described above) was expressed and purified following a modified version of a previously published protocol (A et al., 2020). Briefly, the construct was transfected into 2 L of Expi293 cells and protein was expressed for 48 h. Cells were harvested by centrifugation at 300 × g for 15 min and resuspended in 100 ml of EB-S buffer (100 mM HEPES, pH 7.4, 500 mM NaCl, 5 mM EDTA, 1 mM DTT, 1% Triton X-100, supplemented with 2 μg/ml leupeptin, 10 μg/ml aprotinin, 2 μg/ml pepstatin A, 1 μg/ml calpeptin, 1 μg/ml calpain inhibitor I, 1 mM benzamidine, and 1:1,000 dilution of universal nuclease [Thermo Fisher; 88702]). Cells were lysed using a microfluidizer. Cell debris was removed by ultracentrifugation at 185,000 × g for 1 h (Ti45 rotor, Beckman). The supernatant was incubated with 20 μg/ml avidin (Sigma-Aldrich; 189725) to block endogenous biotin before loading onto a 5 ml Strep-Tactin Superflow column (IBA; 2-1206-025) pre-equilibrated with EB-S buffer. The column was washed extensively with WB buffer (10 mM HEPES, pH 7.4, 150 mM NaCl, 1 mM EDTA, 1 mM DTT) and eluted with Strep elution buffer (10 mM HEPES, pH 7.4, 150 mM NaCl, 1 mM EDTA, 1 mM DTT, and 2.5 mM desthiobiotin). Eluted protein was concentrated using a 100 kDa MWCO Amicon Ultra-15 centrifugal filter (Millipore) and further purified on a Superdex 200 Increase 10/300 GL column (GE Biosciences) equilibrated in 1x K50MEHD buffer (50 mM KCl, 1 mM MgCl_2_, 1 mM EGTA, 10 mM HEPES, pH 7.4, and 1 mM DTT). Protein was aliquoted and snap-frozen in liquid nitrogen.

INF2-FH1-FH2-C (FFC) constructs (wild-type and the chimera with mDia1-DAD) and INF2 1–420 constructs (wild-type, W11L14L18>A mutant, ΔNT, and LM), and the INF2 C-terminal region (CT) were expressed in *E. coli* as GST fusion proteins, as described previously (Gaillard et al., 2011). Bacterial lysates were applied to glutathione-Sepharose resin (GE Biosciences; 17513201), and bound proteins were cleaved on-resin with either tobacco etch virus (TEV) protease for INF2-FFC or thrombin for INF2 1– 420 and INF2-CT. Proteins were further purified by gel filtration using a Superdex 200 16/600 column (GE Biosciences) equilibrated with 1x K50MEHD buffer.

Profilin I was expressed in E. coli and purified as described previously (Harris et al., 2004). Thymosin β4 was also expressed in E. coli in the same manner, resuspended in buffer containing 10 mM HEPES, pH 7.4, and 1 mM EDTA, and lysed by sonication. Lysates were cleared by ultracentrifugation at 185,000 × g for 1 h at 4°C (Ti45 rotor, Beckman), and the supernatant was subjected to stepwise ammonium sulfate precipitation (40%, 60%, and 80% saturation). The 80% supernatant was dialyzed into water (three 24 hr dialyses in 20 L), then lyophilized in a speed vac. Final protein preparations were aliquoted and snap-frozen in liquid nitrogen.

For INF2 constructs, concentration of the final fraction was determined by Bradford assay (BioRad 5000006) and confirmed by Coomassie-stained SDS-PAGE, using known amounts of actin as standards. For profilin and thymosin, concentration and purity of the final fraction was determined by amino acid analysis (Molecular Structure Facility, UC Davis).

### Preparation of INF2-N terminal FITC labeled peptide

N-terminally FITC-Ahx–labeled INF2 N-terminal peptide (FITC-MSVKEGAQRKWAALKEKLGPQDSDPTEAN) was synthesized by BIOMATIK. A total of 5 mg peptide was dissolved in Milli-Q water to 1 mM, and the pH was adjusted to 7.0 using NaOH.

### Fluorescent labeling of INF2-C term

INF2-Cterm was dialyzed overnight against 1 L of DB buffer (2 mM NaPO4, pH 7.0; 100 mM NaCl; 0.5 mM EDTA; 0.25 mM DTT). A 37.5 µM solution of the protein was then mixed with 375 µM fluorescein succinimide (FITC) (Thermo Fisher; 46409) in 50 mM NaPO4, pH 6.0, 100 mM NaCl at 25 °C. The labeling reaction was terminated by adding Tris-HCl (pH 8.0) to a final concentration of 100 mM and DTT to 20 mM. The mixture was subsequently gel-filtered using a Superose 12 column (GE Biosciences) equilibrated in polymerization buffer (1× K50MEH). Final protein concentration was measured by Bradford assay, and FITC concentration was determined using an extinction coefficient of 75,000 M⁻¹ cm⁻¹ at 490 nm. The calculated FITC:INF2-Cterm labeling ratio was 1.02.

### mDia1 DAD and INF2-mDia1-DAD chimera

The DAD sequence from mouse mDia1 (DETGVMDSLLEALQSGAAFRRKRG) was used in these experiments. For fluorescence anisotropy binding studies, the peptide was synthesized as described in (Li & Higgs, 2005), and labeled with TAMRA-maleimide following the same procedure as described in that publication. The chimeric INF2-mDia1-DAD constructs were made in both human INF2-FFC and full-length INF2-CAAX by replacing amino acids 968-986 (EEVCVIDALLADIRKGFQL) with the mDia1 sequence above. To generate INF2-mDia1-DAD, bases 2026-3062 (containing amino acids 675-1020) of human INF2 were synthesized (Integrated DNA Technologies) with mDia1-DAD replacing the INF2-DAD. The construct includes an AflII site at the 5’ end and an SbfI site at the 3’ end, that are unique within the human INF2 sequence. The gene fragment was cloned into either GFP-INF2-CAAX or INF2-FH1-FH2-C using the AflII and SbfI restriction sites.

### Cell treatments

For ionomycin or histamine treatment, 5 x 10^5^ cells were plated onto 35-mm glass-bottom dishes (MatTek). The following day, regular growth medium was replaced by 1 ml of live-cell medium (DMEM with 25 mM D-glucose, 4 mM L-glutamine, and 25 mM HEPES, supplemented with 10% newborn calf serum) (Gibco; 31053028). Another 1 ml of live-cell medium containing 8 µM ionomycin (Sigma; I0634; from a 1 mM stock in DMSO) or 100 µM histaimine (Sigma; H7125; from a 100 mM stock in DMSO) was added swiftly in a circular motion onto the outside of the plate (so as not to activate CIA by mechanical stimulation, (Shao et al., 2015). For ionomycin treatment during live-cell imaging, ionomycin stimulation was applied at the fifth frame (15-second intervals; four imaging fields selected) and imaging was continued for an additional 5 minutes. For histamine stimulation, the procedure was the same, except images were acquired at 10-second intervals for an additional 3 minutes and 20 seconds after drug addition. For fixed-cell imaging, medium was removed and cells were fixed in pre-warmed 4% paraformaldehyde (PFA; 15170; Electron Microscopy Sciences) and 0.25% glutaraldehyde (16020; Electron Microscopy Sciences) in PBS for 15 minutes at room temperature. After fixation, cells were washed three times with PBS and permeabilized with 0.1% Triton X-100 in PBS for 1 minute. Cells were then washed again with PBS and blocked with 10% calf serum in PBS for 20 minutes. Actin filaments were stained with 1 µM TRITC–phalloidin (Sigma-Aldrich, P1951) and DNA was stained with 0.1 mg/L (w/v) 4,6-diamidino-2-phenylindole (DAPI; Calbiochem, 268298). Cells were washed three times with PBS and stored in PBS at 4 °C until imaging.

### Microscopy

Both fixed and live cell samples were imaged using the Dragonfly 302 spinning disk confocal system (Andor Technology) mounted on a Nikon Ti-E base, equipped with an iXon Ultra 888 EMCCD camera and a Tokai Hit stage-top incubator maintained at 37°C. A solid-state 405 smart diode 100-mW laser, solid-state 488 OPSL smart laser 50-mW laser, and a solid-state 560 OPSL smart laser 50-mW laser were used for excitation. Images were acquired with a 100× 1.4 NA CFI Plan Apo objective (Nikon) using Fusion software (Andor Technology).

### CIA intensity measurement in live and fixed cells

For both live-cell and fixed-cell imaging, exposure times and laser intensities were kept constant across all cell lines and treatment conditions. A 25 µm² circular region was selected in the perinuclear cytoplasmic area of each cell, specifically within a medial confocal plane to avoid signal contributions from basal stress fibers or cortical actin. Mean fluorescence intensity (arbitrary units) in this region was measured using ImageJ. For live-cell imaging, Ftractin values at each time point after drug treatment (F) were normalized to the baseline fluorescence (F₀; average of the first four frames) and plotted over time as F/F₀ using Microsoft Excel. For fixed-cell imaging, TRITC–phalloidin fluorescence intensity values were normalized to the TRITC fluorescence measured from the corresponding nuclear area in each cell and plotted as dot plot using GraphPad Prism. Statistical significance between two groups was determined using unpaired Student’s t-tests, comparing the full datasets as indicated in the figure legends.

### Actin preparation

Rabbit skeletal muscle actin was purified from acetone powder as previously described (Spudich and Watt, 1971) and labeled with either pyrenyliodoacetamide (Molecular Probes; P29) or TAMRA NHS ester (Molecular Probes; C1171). Both unlabeled and labeled actin were gel-filtered using a Superdex 75 column and stored in dialysis in G-buffer (2 mM Tris-HCl, pH 8.0, 0.5 mM DTT, 0.2 mM ATP, 0.1 mM CaCl₂, and 0.01% sodium azide) at 4 °C. Dialysis buffer changed weekly.

### Low speed pellet (LSP) preparation

Cells were plated at 4 × 10^6^ cells per 100 mm dish, with two dishes prepared for each cell line. The following day, cells were washed twice with 5 ml PBS and collected using a cell scraper in 2 ml of 2× extraction buffer (1× KMEH: 50 mM KCl, 1 mM MgCl2, 1 mM EGTA, 10 mM HEPES, pH 7.4) supplemented with 1 mM DTT and protease inhibitors (leupeptin 2 µg/ml, aprotinin 10 µg/ml, pepstatin A 2 µg/ml, benzamidine 2 mM, calpain inhibitor I (ALLN) 1 µg/ml, calpeptin 1 µg/ml, cathepsin B inhibitor II 0.05 µg/ml). Cells were lysed using a metal Dounce homogenizer (DWK Life Sciences). The total cell lysate was centrifuged at 600 × g (1800 rpm in Beckman Allegra 6R swinging bucket centrifuge) for 5 min at 4°C. The supernatant was removed, and the pellet was washed with 2 ml of 2× extraction buffer, followed by another centrifugation at 600 × g for 5 min at 4°C. The pellet was then resuspended in 0.5 ml of 2× extraction buffer. Total protein concentration in the LSP fraction was determined by Bradford assay (Bio-Rad). For LSP preparation from transiently transfected cells, 2 × 10^6^ cells were plated on a 100 mm dish. The following day, transfections were performed in OPTI-MEM medium (Life Technologies; 31985062) using 7 µg DNA and 10 µl Lipofectamine 2000 (Invitrogen; 11668) per dish for 3.5 hours. LSP was then prepared as described above, with the final resuspension in 200 µl of 2× extraction buffer. For LSP preparation from CAP1 and CAP2 knockdown cells, 1.5 × 10^6^ cells were used. Knockdown was performed as described, and cells were harvested for LSP preparation 48 hours post-transfection. LSP protein concentration determined by BCA assay (ThermoFisher 23250).

### Free calcium concentration calculations

Free calcium was concentrated for our standard actin polymerization buffer (1× KMEH: 50 mM KCl, 1 mM MgCl2, 1 mM EGTA, 10 mM HEPES, pH 7.4) using Ca-EGTA Calculator v1.3 (https://somapp.ucdmc.ucdavis.edu/pharmacology/bers/maxchelator/CaEGTA-TS.htm). As an example, total CaCl_2_ added to conduct the free calcium concentration curve in Figure 3C were: 0.113 mM (10 nM free), 0.273 mM (25 nM), 0.463 mM (50 nM free), 0.683 mM (100 nM), 0.893 mM (200 nM free), 0.993 mM (300 nM), 1.048 mM (400 nM free), 1.087 mM (500 nM), and 1.174 mM (1 μM free).

### Pyrene actin polymerization assay and fluorescence anisotropy

Actin monomers in G-buffer (6.7 μM actin, 5% pyrene-labeled) were converted to the Mg²⁺-bound form by adding 0.1 volumes of 10 mM EGTA, 1 mM MgCl_2_ for 2 minutes at 23 °C immediately prior to polymerization. Formin protein or 10 μg of LSP was premixed with other proteins (profilin, thymosin, capping protein) in 1.43× polymerization buffer (71.4 mM KCl, 1.43 mM MgCl₂, 1.43 mM EGTA, 14.3 mM HEPES pH 7.4, 1.9 mM DTT, 1.9 mM Tris-HCl, 0.19 mM ATP, 0.09 mM CaCl₂, and 0.009% w/v NaN₃). Pyrene fluorescence (excitation 365 nm, emission 410 nm) was measured using a 96-well fluorescence plate reader (Infinite M1000; Tecan), starting within 1 minute of mixing actin with the formin/LSP premix in the plate. The slope of the fluorescence curve at half-maximum intensity was calculated to represent INF2 activity using KaleidaGraph 4.5 (Synergy Software).

Anisotropy readings were conducted with FITC- or TAMRA-labeled peptides and the indicated binding proteins, in 50 mM KCl, 1 mM MgCl2, 1 mM EGTA, 10 mM Hepes pH 7.4, 0.01% Thesit (Sigma-Aldrich).

### Microscopy-based assays using LSP

For TAMRA-actin imaging, LSP was prepared as described above from U2-OS INF2 KO cells or U2-OS INf2 KO stably expressing GFP-INF2-CAAX. Then, 10 μg of LSP was premixed with other proteins (profilin, capping protein) in 1.43× polymerization buffer (71.4 mM KCl, 1.43 mM MgCl₂, 1.43 mM EGTA, 14.3 mM HEPES pH 7.4, 1.9 mM DTT, 1.9 mM Tris-HCl, 0.19 mM ATP, 0.09 mM CaCl₂, and 0.009% w/v NaN₃). Actin monomers in G-buffer (6.7 μM actin, 10% TAMRA-labeled) were converted to the Mg²⁺-bound form by adding 1 mM EGTA and 0.1 mM MgCl₂ (from a 10× stock) for 2 minutes at 23 °C immediately prior to polymerization. Actin was then mixed with LSP and other components, incubated at 23 °C for 10 minutes, and subsequently placed on ice. The mixture was then combined with an equal volume of 1× extraction buffer containing 0.2 mg/L (w/v) DAPI (Calbiochem, 268298). A 2 μl aliquot was mounted onto a glass slide and covered with a coverslip (72222-01; Electron Microscopy Sciences) for imaging. For TRITC-phalloidin imaging, 10 μg of LSP was incubated with 1 μM TRITC–phalloidin and 0.1 mg/L (w/v) DAPI in 100 μl of 1× extraction buffer. Samples were imaged after a 10-minute incubation at room temperature.

### Calculation of ratios of actin and actin binding proteins in U2OS cells

We used publically-available data from the DepMap portal (Broad Institute, 25Q2 dataset) to determine overall transcript levels for actin, capping protein, thymosin, and profilin. Values used are in TPM (transcripts per million). For actin, thymosin, and profilin, we used the sum of expression from all genes. Actin transcript levels (seven genes): ACTA1, 0.11; ACTA2, 2.40; ACTB, 3843.33; ACTBL2, 3.56; ACTC1, 0.10; ACTG1, 1425.99; ACTG2, 0.24; total actin, 5275.73. Thymosin transcript levels (five genes): TMSB4X, 3236.75; TMSB4Y, 0.00; TMSB10, 2973.20; TMSB15A, 108.11; TMSB15B, 14.45; TMSB15C, 3.99; total thymosin, 6336.50. Profilin transcript levels (four genes): PFN1, 1243.73; PFN2, 163.93; PFN3, 0.00; PFN4, 1.11; total profilin, 1408.77. For the capping protein heterodimer, we used the average of CAPZA (sum of the three CAPZA genes) and CAPZB (single gene). Capping protein transcript levels (four genes): CAPZA1, 138.76; CAPZA2, 51.42; CAPZA3, 0.00; CAPZB, 168.44; total capping protein, 179.31. From these values, the ratios of actin:capping protein:thymosin:profilin are 1:0.034:1.20:0.27.

### Actin pelleting assay

Performed as previously described (Kage et al., 2022) with modifications. Briefly, U2-OS cells were seeded 6 x 10^5^ cells per 35 mm well the day prior to the experiment. The following day, cells were treated with one of three conditions: live cell medium alone, 4 µM ionomycin in live-cell medium for the indicated time or 100 µM histamine in U2-OS culture medium for the indicated time. Following treatment, the medium was removed quickly, and cells were extracted with 2ml of extraction buffer (1xNa50MEH [500 mM NaCl, 20 mM MgCl2, 5 mM EGTA, 100 mM HEPES], 1 mM DTT, 0.4 µM phalloidin, 0.4 µM LatA, and 1% Triton X-100). From each extract, 750 µl was collected as the input sample and mixed with 250 µl of 4x sample buffer (500 mM Tris, pH 6.8, 4 mM EDTA, 40% glycerol, 8% SDS, 40 mM DTT). An additional 1 ml of lysate was transferred to a TLA100 ultracentrifuge tube and centrifuged at 80,000 rpm in a TLA120 rotor (Beckman) for 22 minutes at 4 °C. After centrifugation, 750 µl of the supernatant (containing monomeric actin) was carefully removed and mixed with 250 µl of 4x sample buffer. The pellet (containing filamentous actin) was washed once with 1 ml of 1× Na50MEH containing 1% Triton X-100 (by carefully adding then immediately removing), then resuspended in 1.33 ml of 1x sample buffer. All samples were heated at 95 °C for 5 minutes. Ten microliters of each sample was loaded onto SDS-PAGE gels alongside standard dilutions of purified actin and tubulin protein. Proteins were analyzed by Western blot using an Odyssey CLx imager (LI-COR), using the following antibodies: anti-actin (mouse, mab1501R; Millipore; 1: 10,000) and anti-tubulin (mouse, T9026; Millipore; 1: 10,000). Intensities of actin and tubulin bands determined using LI-COR software Image Studio Lite Ver. 5.2.

For the actin pelleting assay of LSP, 150 µg of LSP was resuspended in 200 µl of extraction buffer containing 1 unit benzonase (Millipore; 70664) along with other additives as indicated (2 µM phalloidin, and 2 µM LatA) and incubated overnight at 4°C. Samples were then centrifuged at 80,000 rpm in a TLA 100 rotor for 22 minutes at 4 °C. After centrifugation, 150 µl of the supernatant (containing monomeric actin) was carefully removed and mixed with 50 µl of 4x sample buffer. The pellet (containing filamentous actin) was washed once with 200 µl of 1× Na50MEH containing 1% Triton X-100, then resuspended in 267 µl of 1x sample buffer.

Calculation of theoretical rate maximal cellular rate of actin addition to INF2-bound barbed ends took into account the on-rate of an actin monomer onto an INF2-bound barbed end in the presence of profilin (∼6.5 mM^-1^sec^-1^, Gurel et al., 2015), 192 million actin molecules/cell in U2-OS (the mean of three independent measurements of total actin content conducted in our laboratory (Hatch et al., 2016; A et al., 2020; Kage et al., 2022), a cytoplasmic volume of 3.14 pL (A et al., 2020), and an estimate that 50% of the cellular actin is monomeric at the time of cell stimulation. Based on these values, U2-OS cells possess 91 mM total actin, of which 45.5 mM is monomeric. Each INF2-bound barbed end should, therefore, allow 296 actins per sec to add to the barbed end. Our estimate of 35 actins/sec/INF2 is lower than this value.

### Actin quantification in LSP

For the quantification of actin in the LSP fraction (Supplementary Figure 2C), U2-OS INF2 KO cells stably expressing GFP-INF2-CAAX were seeded at 4 x 10^6^ cells per 100 mm dish, with two dishes prepared. LSP was prepared as previously described. Briefly, cells were extracted with 2.2 ml of extraction buffer. A 0.2 ml aliquot of the total cell extract was reserved, and the remaining 2 ml was processed for LSP preparation. The resulting LSP was resuspended in 0.5 ml of extraction buffer. Protein concentrations of both the total cell extract and the LSP were determined using the Bradford assay, and the indicated amount of proteins were loaded onto SDS-PAGE gel alongside actin and GAPDH protein standards. Proteins were analyzed by standard Western blot using using the following antibodies: anti-actin (mouse, mab1501R; Millipore; 1: 10,000) and anti-GAPDH (mouse, sc-365062 (G-9); Santa Cruz; 1: 3,000).

### Computational Modeling

A structural model of full-length human INF2-CAAX (Uniprot Q27J81-1) was made using AlphaFold3. AlphaFold returned five models, of which the one with the highest confidence score is depicted in Supplementary Figure 8, though the orientations of DID, DAD, and N-terminus were similar in the other four models produced. A second structural model of a human INF2 variant lacking the DAD (https://alphafold.ebi.ac.uk/entry/A0A087X118) was also used. This model was further subjected to molecular dynamics simulations using Amber 22. The model was prepared for simulation using the s19SB force field (Tian et al., 2020) and solvated in an octagonal box of OPC3 waters (Izadi and Onufriev, 2016) with a minimum of 8 Å between the protein and the edge of the box in all dimensions. Hydrogen mass repartitioning (Hopkins et al., 2015) was applied to allow for a 4-fs simulation time step. The model was energy minimized over 5000 steps, then heated at constant volume from 200 to 310 K over 11,000 4-fs steps. Finally, 100-ns of constant-pressure simulation were performed. Simulations used the Andersen thermostat (DOI: <u>10.1063/1.439486</u>) with a frequency of 1000 steps.

### Data processing and statistical analyses

For the cell imaging data, brightness and contrast were adjusted uniformly using ImageJ software. Figures were compiled and finalized using PowerPoint 2016. Data processing was performed in ImageJ and Microsoft Excel. Statistical analyses, including p-value calculations, were conducted using GraphPad Prism version 6.01. Comparisons between two groups were made using unpaired Student’s *t*-tests. Statistical significance was defined as *p* ≤ 0.05 and is indicated by an asterisk in figure panels. Additional significance levels are represented as follows: ** for *p* ≤ 0.01, *** for *p* ≤ 0.001, and **** for *p* ≤ 0.0001.

## Supplementary figure legends

**Supplementary figure 1.**
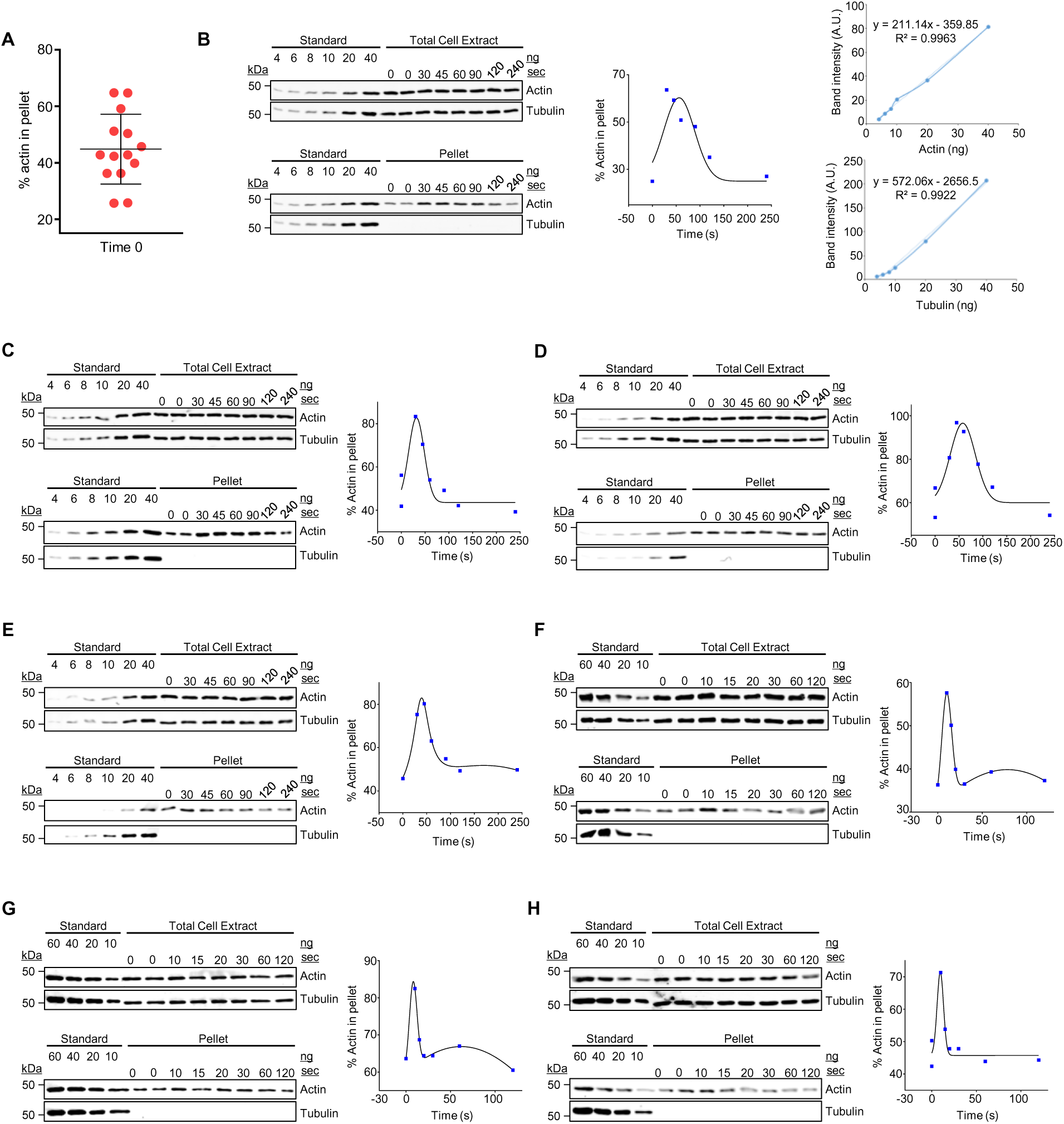
Quantification of pelleted actin in U2-OS cells following ionomycin or histamine treatment. **(A)** Plot showing the percentage of total actin that is pelleted in un-stimulatedU2-OS cells. Data from 7 independent experiments (13 data points). **(B-H)** Individual data from four independent experiments of ionomycin stimulated cells and three independent experiments of histamine stimulated cells shown here. **(B)** Experiment 1. *Left*: Representative western blots of total cell lysates and pellet fractions from U2-OS cells treated with ionomycin for the indicated times (sec). The pellet blot is the same as in Figure 1B, with the difference being that all the standards are shown. *Middle:* Plot of % actin in the pellet versus stimulation time. *Right:* standard curves for actin and tubulin from cell extract blots. **(C), (D)** and **(E)** Western blots and time course plots for Experiments 2, 3 and 4 of ionomycin stimulated cells. **(F), (G)** and **(H)** Western blots and time course plots for Experiment 1, 2 and 3 of histamine stimulated cells.

**Supplementary figure 2.**
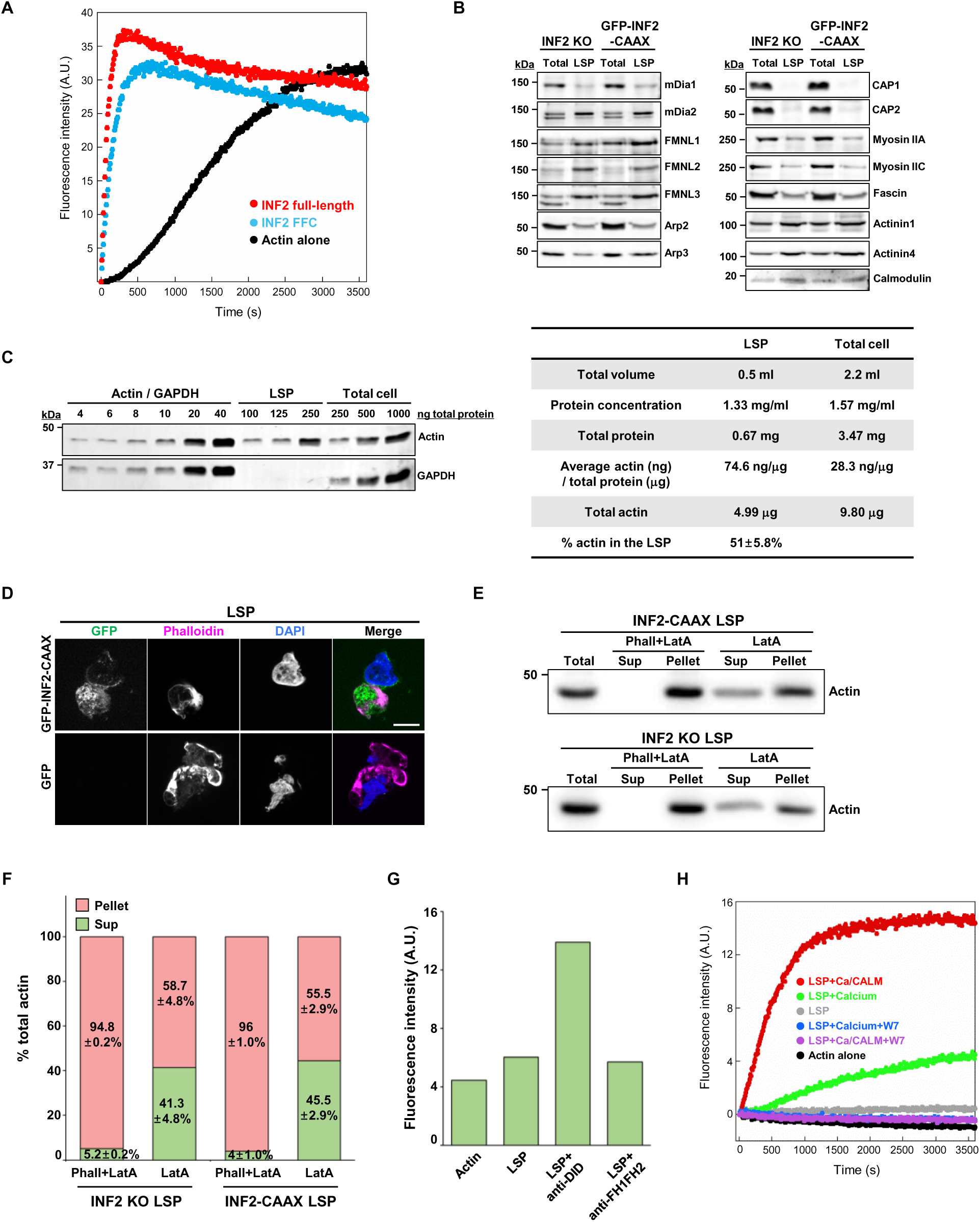
Characterization of cell-free assay. **(A)** Pyrene actin polymerization assays containing 2 µM actin (5% pyrene-labeled), 10 nM INF2 full-length or 10 nM INF2-FFC protein. Experiments were performed in triplicate with similar results, and the data shown are a representative experiment. **(B)** Extended western blots corresponding to Figure 2D. Equal total protein (5 μg) from total cell lysate or LSP of either U2-OS INF2 KO cells or GFP-INF2-CAAX stable cells was loaded and blotted with the indicated antibodies. **(C)** Left: Western blots detecting actin and GAPDH in LSP and total cell lysates from U2-OS INF2 KO cells stably expressing GFP-INF2-CAAX. Right: Table showing the calculated amounts of actin in LSP and total cell lysates. See *Materials and Methods* for details. **(D)** LSPs prepared from either U2-OS INF2 KO cells or cells stably expressing GFP-INF2-CAAX were stained with TRITC-phalloidin and DAPI to visualize filamentous actin associated with LSPs. Scale bar: 10 μm. **(E)** Representative western blots showing actin in the supernatant (Sup) and pellet fractions of LSPs from the indicated cells, LSP treatment with either 2 μM phalloidin + 2 μM LatA, or 2 μM LatA alone overnight. **(F)** The percentage of actin in the supernatant and pellet fractions in LSPs prepared with or without phalloidin. Experiment was performed twice. **(G)** Extended data for figure 2E. Pyrene fluorescence intensity of actin polymerization reactions was measured following overnight incubation. Reactions were performed using 2 μM actin (5% pyrene-labeled), 50 nM capping protein, and 6 μM profilin, with or without 10 μg of LSP from U2-OS INF2 KO cells stably expressing GFP-INF2-CAAX wild-type. Where indicated, 167 nM anti-DID or 167 nM anti-FH1FH2 antibody was included. **(H)** Pyrene-actin polymerization assay containing 2 μM actin (5% pyrene-labeled), 50 nM capping protein, 6 μM profilin, and 10 μg LSP from GFP-INF2-CAAX stable cells, with or without 1 μM calcium, 100 nM CALM and 100 μM W-7 (calmodulin inhibitor). Experiments were performed twice with similar results, and the data shown are a representative experiment

**Supplementary figure 3.**
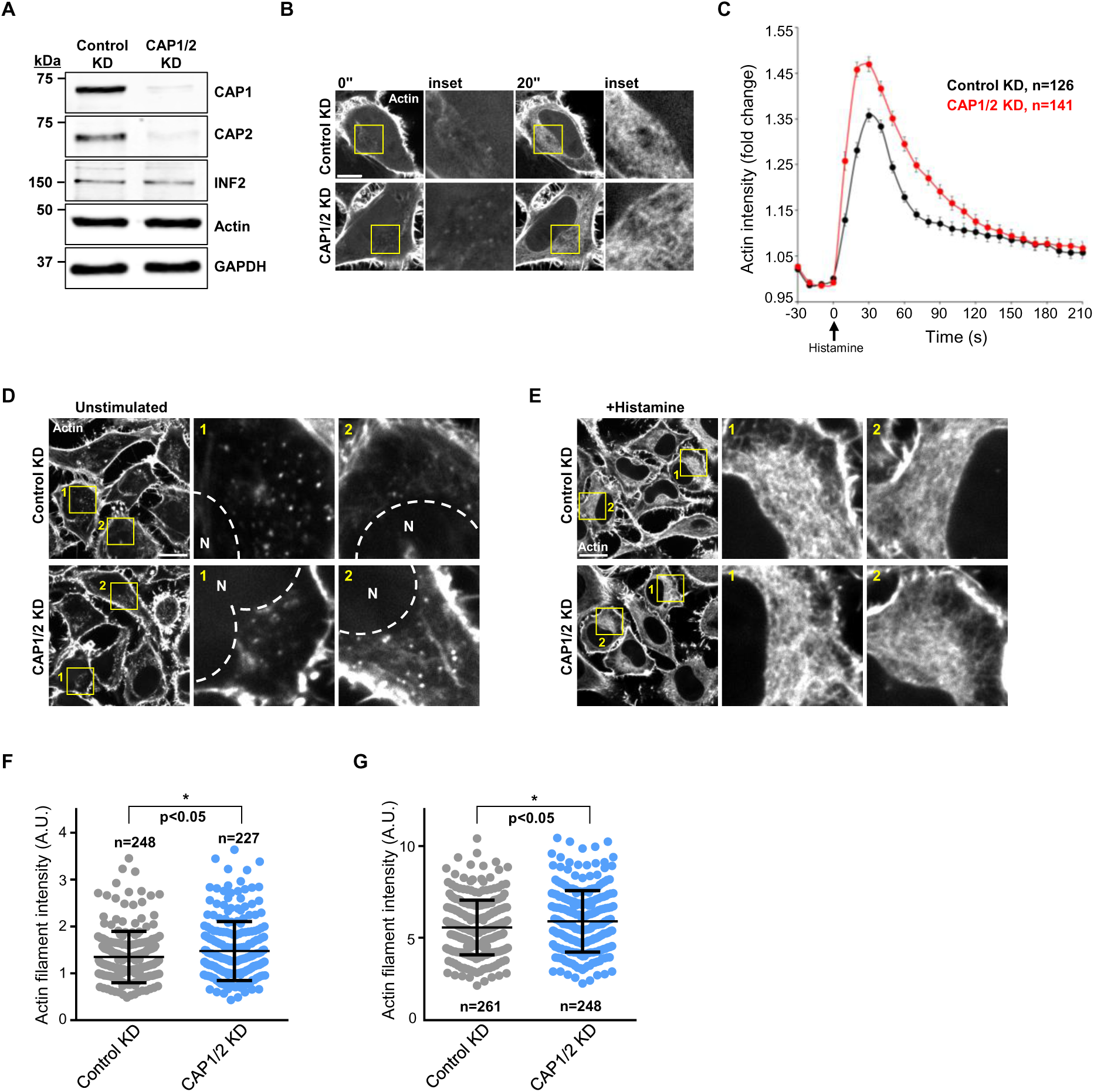
CAP proteins do not influence INF2 regulation in HeLa. **(A)** Western blots of total cell lysate from HeLa cells transfected with either control siRNA or CAP1/2 siRNAs. Equal cell number (10,000) was loaded in each lane and blotted with the indicated antibodies. **(B)** Time-lapse montage of HeLa cells transfected with either control siRNA or CAP1/2 siRNAs, further transfected with mApple-F-tractin, and stimulated with 100 μM histamine at time 0. Insets show magnified views of boxed regions. Scale bar: 10 μm; boxed region: 12 × 12 μm. **(C)** Time-course plot showing changes in cytoplasmic actin intensity (mApple-F-tractin) following histamine addition in HeLa control KD cells and CAP1/2 KD cells. Y-axis values represent fluorescence intensity at each time point normalized to time 0 (F/F₀). Error bars represent the standard error of the mean (SEM), of the stated number of cells analyzed. **(D, E)** HeLa cells transfected with either control siRNA or CAP1/2 siRNA were fixed and stained with TRITC-phalloidin without stimulation (D), or with 100 μM histamine treatment for 30 seconds (E). Insets show magnified views of the boxed regions, as indicated by the corresponding numbers. *N*=nucleus. Scale bar: 20 μm; boxed region: 20 × 20 μm. **(F, G)** Quantification of actin filament intensity in the perinuclear region without stimulation (F) or with 100 μM histamine treatment for 30 seconds (G), normalized to nuclear intensity. Each point represents one region of interest (ROI) per cell. *P*-value was calculated from an unpaired *t-*test. Error bars represent the standard error of the mean (SEM). Experiments were performed twice.

**Supplementary figure 4.**
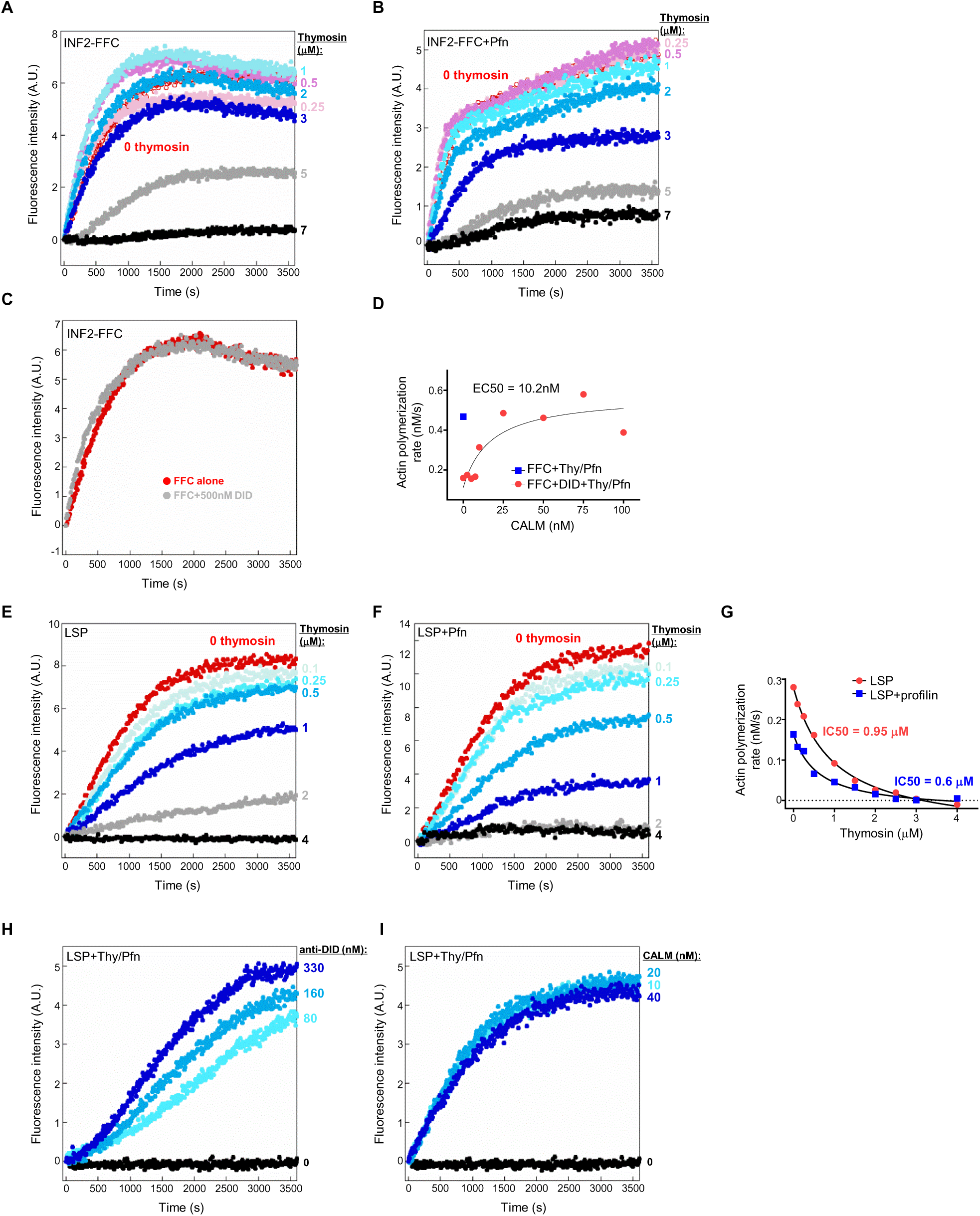
Regulation of INF2-mediated actin polymerization by actin monomer binding proteins. Extended data for figure 6. **(A-C)** Pyrene-actin polymerization assays containing 2 µM actin (5% pyrene-labeled), 50 nM capping protein, and 10 nM INF2-FFC were performed with the following conditions: varying concentrations of thymosin alone (A), varying thymosin plus 2 µM profilin (B), or with/without INF2-DID (C). All experiments were performed in triplicate with similar results, and the data shown are a representative experiment. **(D)** Actin polymerization rates from INF2-FFC inhibition assays in the presence of 2.9 μM thymosin, 6 μM profilin, 50 nM INF2-DID, and varying CALM (1 μM free calcium). **(E-G)** Pyrene-actin polymerization assays containing 2 μM actin (5% pyrene-labeled), 50 nM capping protein, and 10 μg of LSP from U2-OS INF2 KO cells stably expressing GFP-INF2-CAAX were conducted with: varying concentrations of thymosin alone (A), or varying thymosin plus 0.25 μM profilin (B). Actin polymerization rates were calculated from the linear phase of the pyrene-actin polymerization curve (C). **(H, I)** Pyrene-actin polymerization assays containing 2 μM actin (5% pyrene-labeled), 50 nM capping protein, 2.5 μM thymosin, 0.25 μM profilin, and 10 μg of LSP from U2-OS INF2 KO cells stably expressing GFP-INF2-CAAX were performed in the presence or absence of: varying concentrations of anti-DID antibody (D) or CALM in the presence of 1 μM free calcium (E). All experiments were performed twice with similar results, and the data shown are representative of all results.

**Supplementary figure 5.**
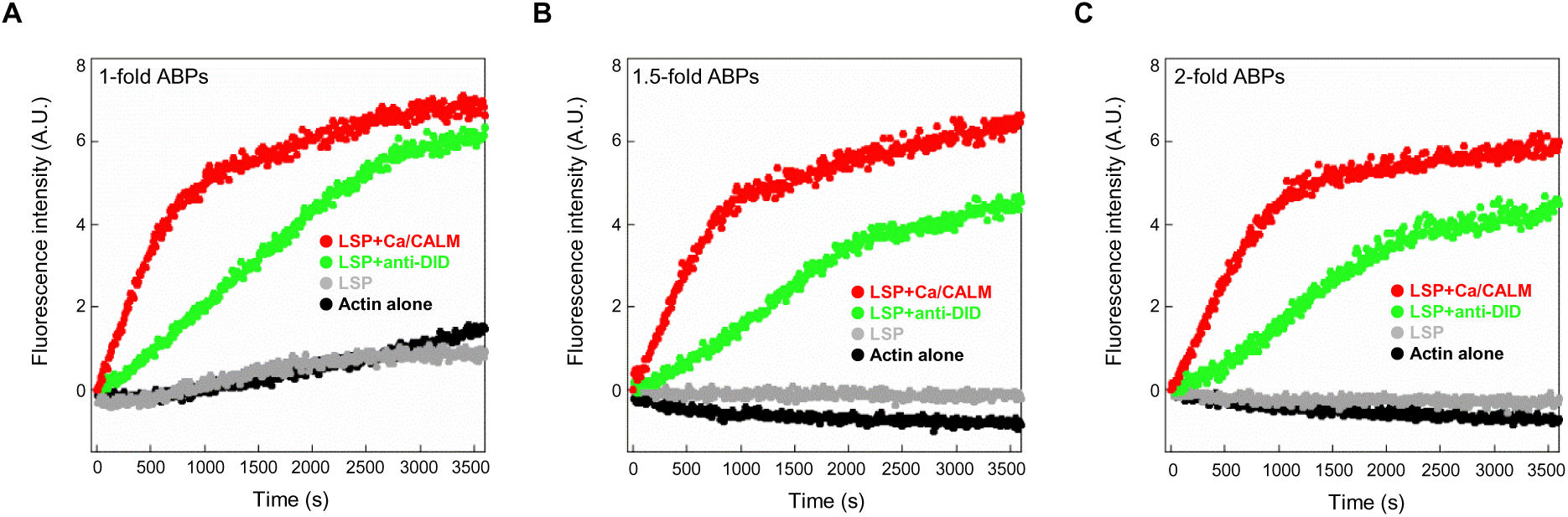
Actin polymerization in the LSP assay using physiologically-relevant ratios of actin:capping protein:thymosin:profilin. Pyrene-actin polymerization assays containing 2 μM actin (5% pyrene-labeled) and 10 μg of LSP from U2-OS INF2 KO cells stably expressing GFP-INF2-CAAX were performed in the presence of **(A)** 68 nM capping protein, 2.4 μM thymosin and 0.53 μM profilin, **(B)** 102 nM capping protein, 3.6 μM thymosin and 0.795 μM profilin, or **(C)** 136 nM capping protein, 4.8 μM thymosin and 1.06 μM profilin, with the indicated components (330 nM anti-DID antibody, or 1 μM calcium with 100 nM CALM). All experiments were performed in triplicate with similar results, and the data shown are representative of all results. See Materials and Methods for calculation of these protein levels.

**Supplementary figure 6.**
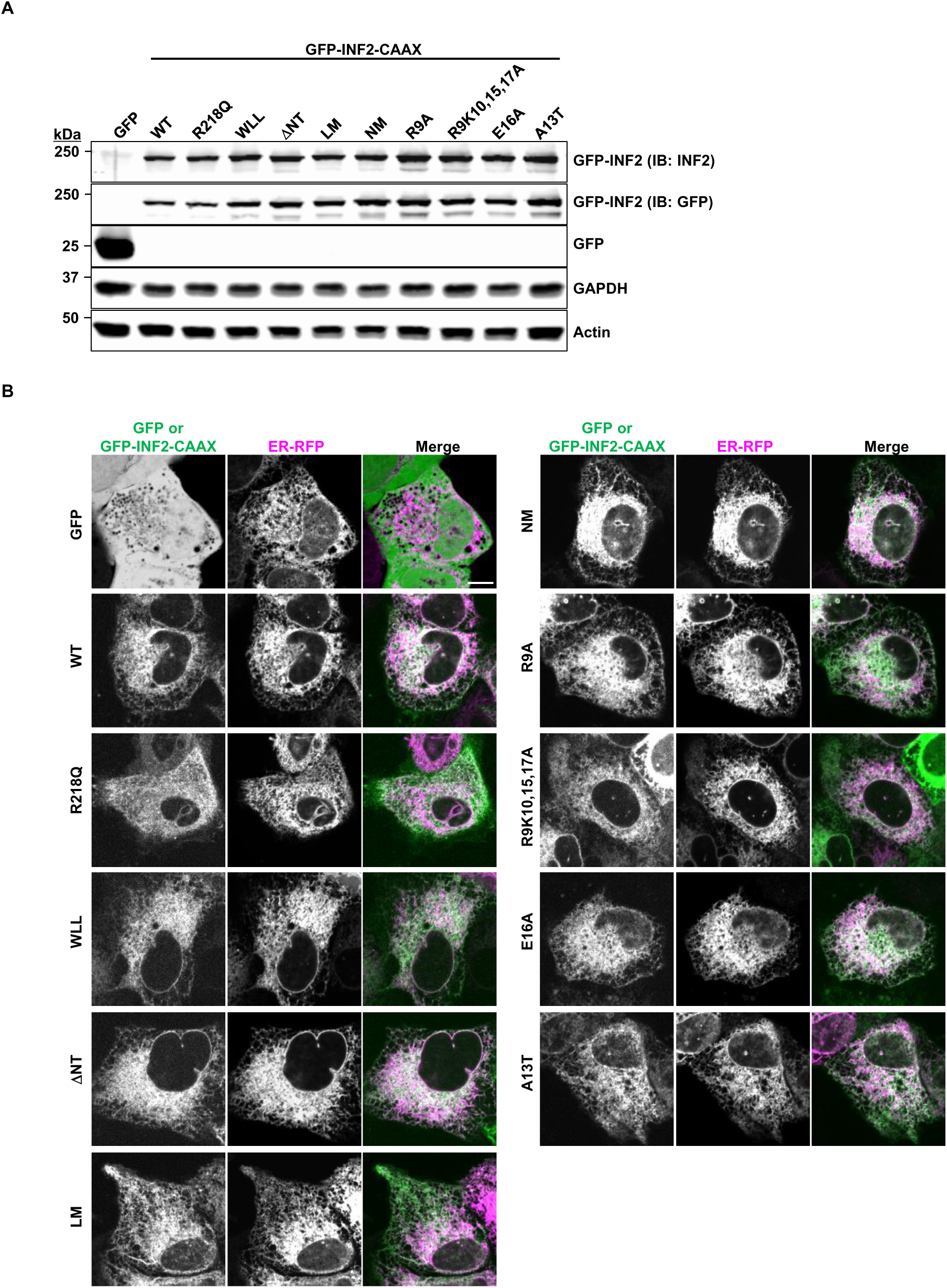
Expression and subcellular localization of GFP-INF2-CAAX wild-type and various N-terminal mutants. **(A)** Western blots of total cell lysate from U2-OS INF2 KO cells transiently expressing the indicated GFP or GFP-INF2-CAAX constructs. Cells were transfected, and total protein was extracted 24 hours post-transfection. (B) Live-cell imaging of U2-OS INF2 KO cells co-transfected with GFP-INF2-CAAX constructs and ER-RFP to assess the subcellular localization of INF2 variants. Scale bar: 10 μm.

**Supplementary figure 7.**
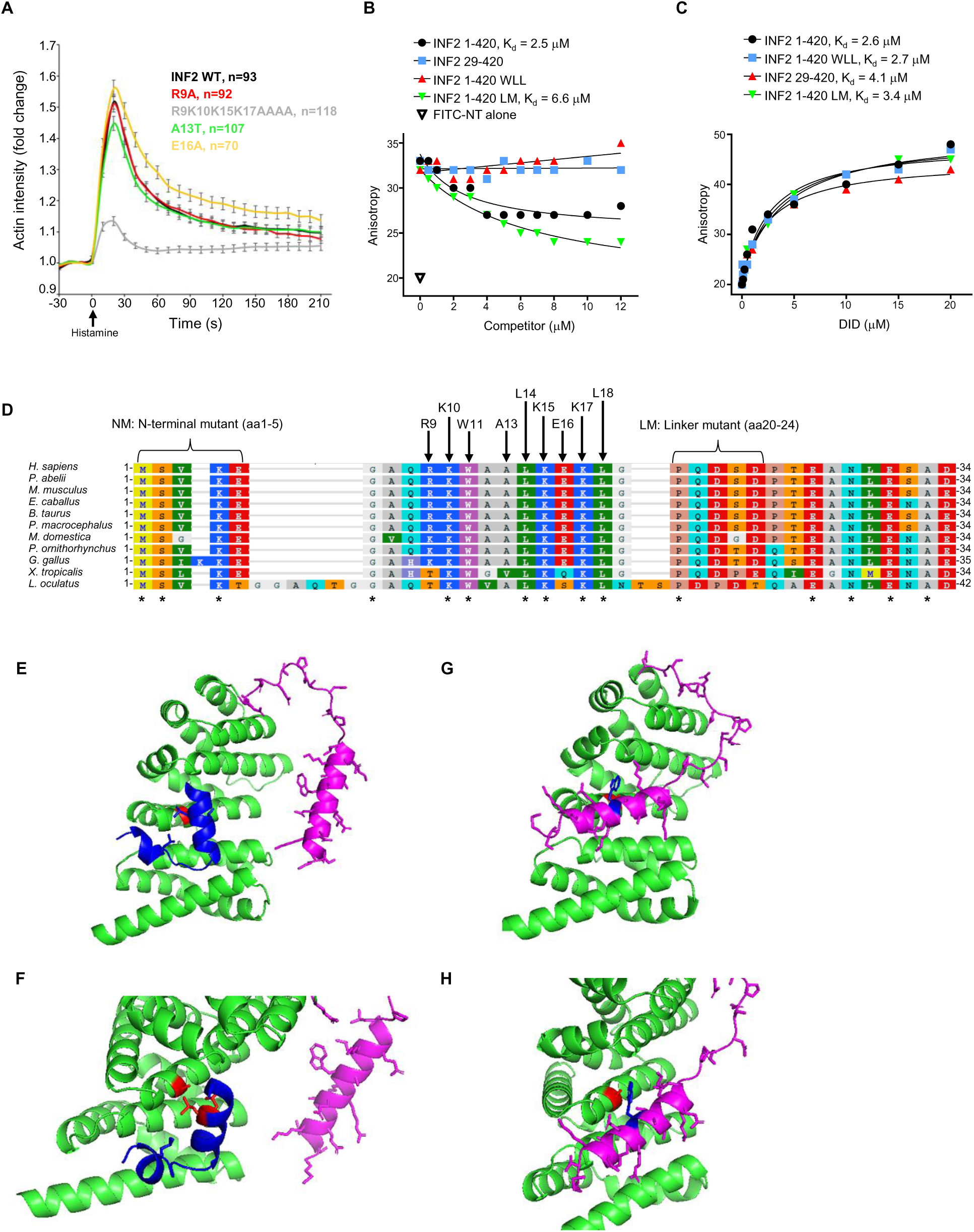
Cellular and structural characterization of INF2 mutants: cellular assays, INF2 DID and INF2 C-terminal interactions, evolutionary conservation, and AlphaFold models of DID interactions. **(A)** Time-course plot showing changes in cytoplasmic actin (mApple-F-tractin) following histamine addition in GFP-INF2-CAAX wild-type or INF2 N-terminal mutants. Y-axis values represent fluorescence intensity at each time point normalized to time 0 (F/F₀). Experiments were performed in triplicate; error bars represent the standard error of the mean (SEM). **(B)** Fluorescence anisotropy competition assays with 100 nM FITC-INF2-NT, 2 μM CALM, 50 μM free calcium, and a range of concentrations of the indicated INF2 construct **(C)** Fluorescence anisotropy assay with 50 nM FITC-INF2 C-terminal region (891-1250 aa) and a range of concentrations of the indicated INF2 DID regions. **(D)** Alignment of INF2 N-terminal sequences across various species. Mutants used in this study are indicated above the aligned sequences. Asterisks (*) indicate identical residues. Sequence alignment was performed using COBALT, with Rasmol color coding for amino acids. Species (with UniProt numbers): H. sapiens (human, Q27J81), P. abelii (orangutan, H2NME2), M. musculus (mouse, Q0GNC1), E. caballus (horse, F6UX87), B. taurus (cow, A0AAA9SIE1), P. macrocephalus (sperm whale, A0A2Y9TAJ8), M. domestica (opossum, F6TXY8), P. ornithorhynchus (platypus, A0A6I8NFG2), G. gallus (chicken, A0A8V0Z7F3), X. tropicalis (frog, A0A6I8SNL0), L. oculatus (fish, W5N4V5). **(E, F)** AlphaFold model of full-length INF2-CAAX, generated in AlphaFold3. Residue A149 in DID and L976 and L977 in DAD shown in red. **(G, H)** AlphaFold model of a human construct lacking the DAD (https://alphafold.ebi.ac.uk/entry/A0A087X118), in which the N-terminal peptide interacts with the DAD-binding region of the DID, albeit in a different orientation. Residue A149 in DID shown in red, and W11 in N-terminus shown in blue. In both models, residues outside of the N-term (purple, 1-34), DID (green, 35-263) and DAD (blue, 972-990) have been omitted. Top: full model. Bottom: zoom to region of interest.

## Acknowledgements

We thank current and past members of the Higgs lab (Rajarshi Chakrabarti, Anna Hatch, Frieda Kage, Sukrut Kamerkar, Will King, Ao Liu, and Ziwei She) for valuable input throughout the project, as well as Luca Cim for activating the process. We are also grateful to William T. Wickner and Gustav E. Lienhard for their thoughtful input and guidance throughout the project. HNH is supported by NIH grants DK088826 and GM122545, as well as P20 GM113132 to BioMT. ML is supported by the National Research Foundation of Korea (NRF), funded by the Ministry of Education (2021R1A6A3A03045359).

